# Individual variability in the relationship between physiological and resting-state fMRI metrics

**DOI:** 10.1101/2024.05.02.592237

**Authors:** Stefano Moia, Gang Chen, Eneko Uruñuela, Rachael C. Stickland, Maite Termenon, César Caballero-Gaudes, Molly G. Bright

## Abstract

Cerebrovascular Reactivity (CVR), the brain’s vascular response to a vasodilatory stimulus, can be measured using fMRI during breathing challenges that modulate arterial CO_2_ levels. CVR is an important indicator of cerebrovascular health, although its estimation can be challenging due to the extra experimental setup and/or the subject compliance required. To overcome these limitations, summary metrics based on resting state fluctuations (RSF), such as the (fractional) amplitude of low frequency fluctuations (f/ALFF) and the resting state fluctuation amplitude (RSFA), have been proposed as alternative estimates of CVR, as they are frequently associated with vascular and physiological factors. Previous studies have reported a significant relationship between CVR estimates obtained by means of respiratory paradigms and RSF metrics. However, the total sample sizes, considering both subjects and sessions, are typically small, and not all studies agree on the degree of the relationship. Furthermore, to our knowledge, these studies have only reported cross-sectional analyses, whereas intra-subject longitudinal relationships between CVR estimates and RSF metrics are unknown. Leveraging a unique dense sampling dataset in which resting state and breath-hold multi-echo fMRI were collected in 7 subjects with 10 sessions each, we provide evidence of high individual variability in the inter-session (i.e. intra-subject) relationship between RSF metrics and CVR. These results indicate that RSF metrics might not be a suitable proxy of CVR in clinical settings or for BOLD signal calibration as they may not properly account for intra-individual physiological variations in BOLD fMRI data.

## 1 Introduction

Cerebrovascular Reactivity (CVR), the ability of vessels to react to vasodilatory and vasoconstrictive stimuli, is an important indicator of cerebrovascular health (Chen & Fierstra, 2021). In recent years, the adoption of Blood Oxygen Level Dependent (BOLD) contrast in functional Magnetic Resonance Imaging (fMRI) has facilitated the study of CVR as a potential biomarker for various pathological conditions (Donahue et al., 2014; Fierstra et al., 2018; Krainik, Hund-Georgiadis, Zysset & Von Cramon, 2005; Marshall et al., 2014; Mikulis et al., 2005).

The gold standard approach to evaluate CVR with BOLD fMRI is based on gas challenges, consisting of subjects inhaling CO_2_ enriched air, often with the aid of gas-delivery systems (for a complete review, see Liu, De Vis & Lu, 2018). However, the extra setup required for gas challenges and the lower tolerance for gas mixtures displayed by certain vulnerable groups, such as patients, children, and elderly people, limits the uptake of such systems (Pinto, Bright, Bulte & Figueiredo, 2021).

Alternatively, respiratory challenges such as breath-holding (BH) have been proposed as a means of modulating vascular CO_2_ levels endogenously (for a complete review, see Pinto et al., 2021). BH challenges have been widely adopted, successfully applied in clinical studies, and can be tolerated better by vulnerable groups (Liu et al., 2017). While the elicited CVR response in BH challenges is comparable to that of inhaled CO_2_ (Kastrup, Krüger, Neumann-Haefelin & Moseley, 2001; Tancredi & Hoge, 2013), the required task compliance can still be considered as a limitation in the adoption of respiratory challenges. Some issues can be overcome, such as subjects performing shorter apnea periods than instructed (providing recordings of end-tidal CO_2_ are available, see Bright & Murphy, 2013), or inadequate recordings of exhaled CO_2_ (Zvolanek et al., 2023). However, more general lack of compliance with task instructions can be particularly problematic, as often seen in clinical settings (Jahanian et al., 2017). For this reason, researchers have looked for alternative approaches for CVR estimation with minimal instructions (e.g. with intermittent breath modulation (Liu et al., 2020)) or, more commonly, directly from fMRI data collected at rest (Hou et al., 2023; Liu et al., 2017; Taneja et al., 2019).

Resting state (RS) fMRI constitutes an attractive alternative to CVR-inducing breathing tasks. Not only is it easier to collect, especially for clinical settings (Golestani, Wei & Chen, 2016), but also multiple RS fMRI datasets are currently available and easy to obtain for reanalysis. Part of the intrinsic (a.k.a. spontaneous) signal fluctuations that contribute to the RS fMRI time series are driven by systemic physiological and vascular factors related to changes in arterial CO_2_ (e.g. see Kannurpatti, Motes, Biswal & Rypma, 2014; Wise, Ide, Poulin & Tracey, 2004). Although the magnitude of such fluctuations depends on the particular acquisition parameters (Bianciardi et al., 2009), such knowledge has been leveraged not only to estimate CVR, but also to normalise BOLD responses to neural factors by taking into account physiological scaling of the signal (Di, Kannurpatti, Rypma & Biswal, 2013; Kalcher et al., 2013; Kannurpatti & Biswal, 2008; Kannurpatti, Motes, Rypma & Biswal, 2011; Kazan et al., 2016; Tsvetanov et al., 2015).

Various methods to retrieve such fluctuations within the RS signal have been proposed, for instance by considering the global signal in particular frequency bands (Liu et al., 2017) or across specific tissue masks (Jahanian et al., 2017; Zerweck et al., 2022), by leveraging the frequency domain rather than the time domain (Xu et al., 2023), by reconstructing CVR maps through deep learning (Hou et al., 2023), or by recursive tracking of slow fluctuations, either correlated to peripheral signals (Tong, Bergethon & Frederick, 2011) or to the global signal itself (Erdoğan, Tong, Hocke, Lindsey & Frederick, 2016). However, the most widespread approach to assess physiological fluctuations in RS probably remains computing voxelwise RS fluctuations (RSF) measures, such as the Resting State Fluctuation Amplitude (RSFA) (Kannurpatti & Biswal, 2008) or its equivalent measure in the frequency domain, the Amplitude of Low Frequency Fluctuations (ALFF) (Zang et al., 2007).

ALFF and RSFA have both been proposed and adopted as low-cost approximations of CVR, due to their significant cross-sectional relationship with induced CVR (Chen et al., 2023; Golestani et al., 2016; Kannurpatti et al., 2014) or the within-subject(s), within-session spatial agreement of the two measures (De Vis, Bhogal, Hendrikse, Petersen & Siero, 2018; Kannurpatti & Biswal, 2008; Kannurpatti et al., 2011; Liu et al., 2013). However, this relationship is a debated topic in the field, due to contrasting evidence from different studies or settings. When such a relationship was observed within-subjects (i.e. spatial correlation), Kannurpatti and Biswal (2008) reported a significant correlation between RSFA, BH-induced CVR, and gas inhalation-induced CVR, particularly in motor areas, while Liu et al. (2013) reported a modest correlation between RSFA and CVR that could indicate that these two measures are not interchangeable (cfr. Lipp, Murphy, Caseras & Wise, 2015). Similarly, in cross-sectional studies Chen et al. (2023) reported significant, but very modest correlation between ALFF and CVR. Likewise, Golestani et al. (2016) reported significant average grey matter agreement between induced CVR and RSFA and ALFF across subjects. However, the latter study also reported poor spatial agreement between CVR, RSFA, and ALFF when assessing this relationship across different regions of interest within the same subjects, showing high variability in the spatial correlation across the examined cohort. Other cross-sectional studies reported less agreement between RSF metrics and induced CVR between-subjects (Lipp et al., 2015), a variable agreement depending on age and task domains (Kannurpatti et al., 2014), or a lower sensitivity to haemodynamic impairment of RSF compared to CVR (De Vis et al., 2018), thereby inviting carefulness in interpreting the relationship between CVR and RSF. Fesharaki et al. (2021) suggest that lower agreement between RSF measures and BH-induced CVR might be attributed to a lack of optimal thresholding of the corresponding maps. More recently, Huck et al. (2023) showed that ALFF has a bias to the properties of venous vasculature, which might confirm the hypothesis that these RSF metrics are biased by resting cerebral blood volume (De Vis et al., 2018), although Bailes, Gomez, Setzer and Lewis (2023) attributes such bias more to the neurovascular coupling than to the CVR mechanism, since the latter does not include the same metabolic factors driving the former (D’Esposito, Deouell & Gazzaley, 2003; Iadecola, 2017; Pinto et al., 2021).

With conflicting reports in the existing literature, and the potential utility of RSF measures for BOLD response normalisation and as a metric of cerebrovascular health in clinical settings, it is important to gain clarity on the relationship between RSF metrics and CVR. In this study, we leverage the first dense mapping dataset of cerebrovascular reactivity mapping, involving 7 subjects undergoing 10 sessions of BOLD fMRI imaging (Moia, Uruñuela, Ferrer & Caballero-Gaudes, 2020), to observe how RSF and BH-induced CVR measures agree across the span of multiple sessions in single subjects (i.e. intra-subject relationship). To our knowledge, this perspective has not yet been explored with BOLD fMRI. We compare CVR maps, optimised for spatial variability in haemodynamic lag (Moia, Stickland et al., 2020) and measured with multi-echo multi-band fMRI to improve its estimation and reliability (Moia et al., 2021), with RSF measures related to physiological fluctuations, i.e. ALFF and RSFA. While the original definition of ALFF included a normalisation of the spatial maps by their average, this last step is often omitted; hence, we report both ALFF and the normalised version of ALFF, mALFF (Zang et al., 2007). As a negative control, we also compare CVR maps to fractional ALFF (Zou et al., 2008), a measure specifically designed to be less susceptible to physiological fluctuations, expecting it to be less related to CVR in comparison to ALFF, mALFF, or RSFA.

## 2 Methods

### 2.1 Dataset

Ten healthy subjects with no record of psychiatric or neurological disorders (5F, age range 24-40 years at the start of the study) underwent ten MRI sessions in a 3T Siemens PrismaFit scanner with a 64-channel head coil. Each session took place one week apart, on the same day of the week and at the same time of the day to minimise effects related to circadian rhythms (see e.g. Shannon et al., 2013). Note that increasing the trial size (in this case, the number of sessions) has a similar impact on statistical efficiency as increasing the number of subjects (Chen et al., 2022).

All participants had to meet several further requirements, i.e. being non-smokers, refraining from smoking for the whole duration of the experiment, and not suffering from respiratory or cardiac health issues. They were also instructed to refrain from consuming caffeinated drinks for two hours before each session. Informed consent was obtained before each session, and the study was approved by the Basque Center on Cognition, Brain and Language ethics committee.

### 2.2 MRI data

Within the MRI session, subjects underwent several runs of multi-echo (ME) fMRI data acquisitions, including a ten-minute RS and an eight-minute BH task that are the focus of this study. This fMRI data was acquired with a T2*-weighted multi-echo simultaneous multi-slice gradientecho echo planar imaging sequence provided by the Center for Magnetic Resonance Research (CMRR, Minnesota) (Moeller et al., 2010; Setsompop et al., 2012). While the number of volume acquisitions was adapted according to the duration of the RS and the BH paradigm (400 and 340 volumes respectively), all other sequence parameters were identical: TR = 1.5 s, TEs = 10.6/28.69/46.78/64.87/82.96 ms, flip angle = 70°, MB acceleration factor = 4, GRAPPA = 2 with Gradient-echo reference scan, 52 slices with interleaved acquisition, Partial-Fourier = 6/8, FoV = 211×211 mm², voxel size = 2.4×2.4×3 mm³, Phase Encoding = AP, bandwidth=2470 Hz/px, LeakBlock kernel reconstruction (Cauley, Polimeni, Bhat, Wald & Setsompop, 2014) and SENSE coil combination (Sotiropoulos et al., 2013). Single-band reference (SBRef) images were also acquired for each TE and each run.

During the fMRI acquisitions, exhaled CO_2_ and O_2_ levels were monitored and recorded (sampling rate: 10 kHz) using a nasal cannula (Intersurgical) connected to an ADInstruments ML206 gas analyser unit and transferred to a BIOPAC MP150 physiological monitoring system. Scan triggers were simultaneously recorded, as well as cardiac pulse data using a photoplethysmograph (PPG, TSD200 transducer with an PPG100C-MRI amplifier), and respiration effort data (using either a TSD221-MRI or TSD110-MRI with a DA100C module, or a TSD201 transducer with an RSP100C amplifier). The physiological recordings started 15 seconds before and lasted at least 15 seconds longer than the ME-fMRI data recordings.

In addition, a pair of Spin Echo EPI images with opposite phase-encoding (AP or PA) directions and identical volume layout (TR = 2920 ms, TE = 28.6 ms, flip angle = 70°) were collected before each functional run in order to be able to estimate field distortions, as suggested in the Human Connectome Project protocol (Glasser et al., 2016).

Subjects also underwent two anatomical images: a T2-weighted Turbo Spin Echo image (TR = 3.39 s, TE = 389 ms, GRAPPA = 2, 176 slices, FoV read = 256 mm, voxel size = 1*×*1*×*1 mm³, TA = 300 s) at the beginning of the MRI session, and a T1-weighted MP2RAGE image (TR = 5 s, TE = 2.98 ms, TI1 = 700 ms, TI2 = 2.5 s„ flip angle 1 = 4°, flip angle 2 = 5°, GRAPPA = 3, 176 slices, FoV read = 256 mm, voxel size = 1*×*1*×*1 mm³, TA = 662 s) at the end of the session.

Note that between the RS and the BH task, subjects underwent another 54 minutes of fMRI data acquisition, consisting of 3 cognitive and sensory tasks interleaved by three RS runs. The choice of using the first RS was dictated by avoiding any possible influence of these tasks in the subsequent RS runs. RS and BH data are available on OpenNeuro (Moia, Uruñuela et al., 2020).

### 2.3 Experimental tasks

Task instructions were explained before the first session, and then briefly reiterated if required before each run took place in every subsequent session. During the RS run, subjects were instructed to remain as still as possible, with their eyes open, while fixating them on a white cross shown in the middle of a black screen through a mirror screen located in the head coil. The BH task was composed of 8 BH trials, with each trial comprised of (a) 24 s of paced breathing with a 6 s period, (b) an end-expiration breathhold of 20 s followed by another exhalation, and (c) 11 s of recovery breathing (Bright & Murphy, 2013; Moia et al., 2021). During the BH task, instructions were provided textually.

Please refer to Moia et al. (2021) for more information on the MRI and physiological recording parameters and data. Note that 3 subjects did not complete all ten BH tasks successfully. Consequently, the dataset used in this study includes seven subjects (4F, age 25-40y) undergoing ten sessions per subject.

### 2.4 Data preprocessing and analysis

All data analysis scripts, based on AFNI (Cox, 1996), ANTs (Tustison et al., 2014), FSL (Jenkinson, Beckmann, Behrens, Woolrich & Smith, 2012), numpy (Harris et al., 2020), scipy (Virtanen et al., 2020), peakdet (Markello & DuPre, 2020), and phys2cvr (Moia, 2022), are publicly available on GitHub^1^. Data preprocessing and analysis was run in a singularity container, also publicly available^2^.

Raw data were reorganised following the Brain Imaging Data Standard (Gorgolewski et al., 2016) using BIDScoin (Zwiers, Moia & Oostenveld, 2022) for the MRI data, and phys2bids (phys2bids developers et al., 2019) for the physiological recordings. Then, the fMRI data of the BH task were preprocessed and analysed following the “Optimal Combination” approach (OC-MPR) described in Moia, Stickland et al. (2020) and Moia et al. (2021), where a lagged General Linear Modelling strategy was applied to the BH fMRI data to obtain a CVR map (in units of %BOLD/mmHg P_ET_CO_2_) optimised for voxelwise haemodynamic delay for each scan session.

Briefly, the SBRef image of the BH run was first skull-stripped and corrected for field susceptibility distortion with the TOPUP approach (Andersson, Skare & Ashburner, 2003) using the two Spin Echo EPI images with opposite phase-encoding directions, which were skull-stripped previously. Next, the T2-weighted image was skull-stripped and co-registered to the MP2RAGE image and to the SBRef image. These spatial transformations were combined in a single one, which was used to transform the brain mask obtained with the T2-weighted image into the space of the T1-weighted MP2RAGE image. A finite mixture modelling segmentation in 3 tissue compartments was run on the masked T1-weighted MP2RAGE image with Atropos (Avants, Tustison, Wu, Cook & Gee, 2011) to obtain a binary mask of grey matter (GM) voxels that was transformed back into the SBRef space.

The first 10 volumes of the BH functional time series were removed to ensure steady-state magnetisation, after which image realignment to the SBRef was estimated using the first echo, and this realignment was then applied to all echoes. Echo series were then linearly combined with T_2_*-dependent weights to obtain the optimally combined (OC) dataset (DuPre et al., 2021; Posse et al., 1999). Finally, the OC volume was corrected for geometric distortions using the same procedure used for the SBRef image.

CO_2_ data were downsampled to 40 Hz and further time-pass filtered to reduce frequencies above 2 Hz (low-pass Butterworth filter of seventh order). Subsequently, end-tidal peaks were automatically and manually classified, linearly interpolated, and convolved with the two-gamma variate canonical hemodynamic response function (HRF) to obtain a P_ET_CO_2_*hrf* trace per run. The P_ET_CO_2_*hrf* trace, optimally combined BH time series, and GM mask were used to compute CVR maps and haemodynamic delay maps using phys2cvr (Moia, 2022). phys2cvr implements a lagged-GLM approach, using a time span of 18 seconds centered around the alignment of the P_ET_CO_2_*hrf* trace and the average GM time series, with a step of .3 seconds. In addition to the shifted P_ET_CO_2_*hrf* trace, the lagged-GLM also comprised the 6 realignment parameters, their 6 temporal derivatives and Legendre polynomials up to 4^th^ order as confounding factors to account for motion-related effects and slow frequency trends in the OC signal (Moia, Stickland et al., 2020; Moia et al., 2021).

RS data underwent the same data preprocessing applied to the BH data, except that the 12 realignment parameters and Legendre polynomials up to 4^th^ order were regressed out of the data after correcting for geometric distortions, since it is not possible to add them to the estimation of RSF metrics. Then, RSFA, ALFF, mALFF, and fALFF were estimated with 3dRSFC (Taylor & Saad, 2013) considering a frequency band between 0.01 and 0.1 Hz.

The average value of CVR and of each RSF metric across GM voxels was computed per subject and session. Finally, to compute voxelwise relationships, all CVR and RSF metric maps were finally normalised to the MNI152 template (packaged with FSL and resampled at 2.5 mm isotropic voxel resolution) and spatially smoothed with a Gaussian kernel with 5 mm FWHM within a dilated GM mask (dilation of 1 voxel, considering only face neighbours).

### 2.5 Relationship between RS metrics and CVR values

#### 2.5.1 Groupwise relationship

A set of Linear Mixed Effects (LME) models was set up using R (Bates, Mächler, Bolker & Walker, 2015; Fox & Weisberg, 2019; Kuznetsova, Brockhoff & Christensen, 2017; Neuwirth, 2022; R Core Team, 2020; Wickham, 2016; Wilke, 2023) for the average GM values and using 3dLMEr (Chen, Saad, Britton, Pine & Cox, 2013) for the voxelwise maps in order to estimate the relationship between CVR and each of the RSF metrics (RSFA, ALFF, mALFF, and fALFF) at the group level:

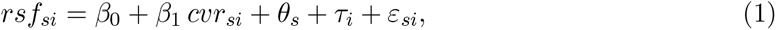

where *rsf_si_* denotes either the RSFA, ALFF, mALFF, or fALFF map of the *i*th participant during the *s*th session, *cvr_si_* denotes the corresponding CVR map, index *s* denotes the session, *i* denotes the subject, *β*_0_ denotes the intercept, *β*_1_ denotes the fixed effect of *cvr_si_* on *rsf_si_*, *θ_s_* and *τ_i_* denote random effects of sessions and individuals, and *ε_si_* denotes noise. Note that all values, both in the voxelwise and in the average GM case, were subject-centered (i.e. demeaned by each subject average) before the coefficients of the LME model were estimated.

As no voxel survived the threshold of *p ≤* 0.05 after controlling for multiple comparisons with the FDR procedure (Benjamini, Krieger & Yekutieli, 2006), the results are reported at an uncorrected significance threshold of *p ≤* 0.05, both thresholded and non-thresholded but modulated by statistical significance.

#### 2.5.2 Subjectwise relationship

A second set of linear models was used to assess the same relationships between RSF metrics and CVR at the subject level using R for average GM values and 3dMVM (Chen, Adleman, Saad, Leibenluft & Cox, 2014) for voxelwise maps.

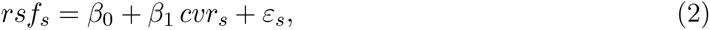

where *rsf_s_* denotes either the RSFA, ALFF, mALFF, or fALFF map during the *s*th session, *cvr_s_* denotes the corresponding CVR map, *β*_1_ denotes the fixed effect of *cvr_s_* on *rsf_s_*, and *ε_s_* denotes noise.

As no voxel survived the threshold of *p ≤* 0.05 after controlling for multiple comparisons with the FDR procedure (Benjamini et al., 2006), the results are reported at an uncorrected significance threshold of *p ≤* 0.05, both thresholded and non-thresholded but modulated by statistical significance.

## 3 Results

Figure 1 shows the CVR, RSFA, ALFF, and fALFF maps of a representative session of a representative subject, as well as the whole brain distribution of CVR, RSFA, ALFF, and fALFF values. Because different metrics exhibit different amplitude ranges, and their amplitude was not rescaled nor normalised, the display ranges presented here are different for each map, but the upper limit is consistently the 99^th^ percentile of each amplitude distribution, considering positive, non-zero voxels only. Note that mALFF is already a rescaled version of ALFF, hence its map is omitted here, but is depicted in Figure S1. Beside the different data range, RSFA and (m)ALFF maps show minimal difference in spatial distribution. Both of them resemble highly the CVR map, in particular regarding the areas presenting the highest positive values across the brain and their value distribution skewness and peak matches closely the positive side of the distribution of CVR. Conversely, and as expected, the fALFF amplitude distribution is qualitatively very different from ALFF and RSFA, appearing flatter than the others and shifted toward higher values, while its map shows more uniform values than the other three, especially within the same cerebral tissues. CVR is the only metric that presents meaningful negative values, that can be interpreted as the effect of a blood flow redistribution ("steal physiology", see Sobczyk et al., 2014; though see Stickland et al., 2021), or as the effect of a reduction of CSF volume (Thomas et al., 2013).

**Figure 1:**
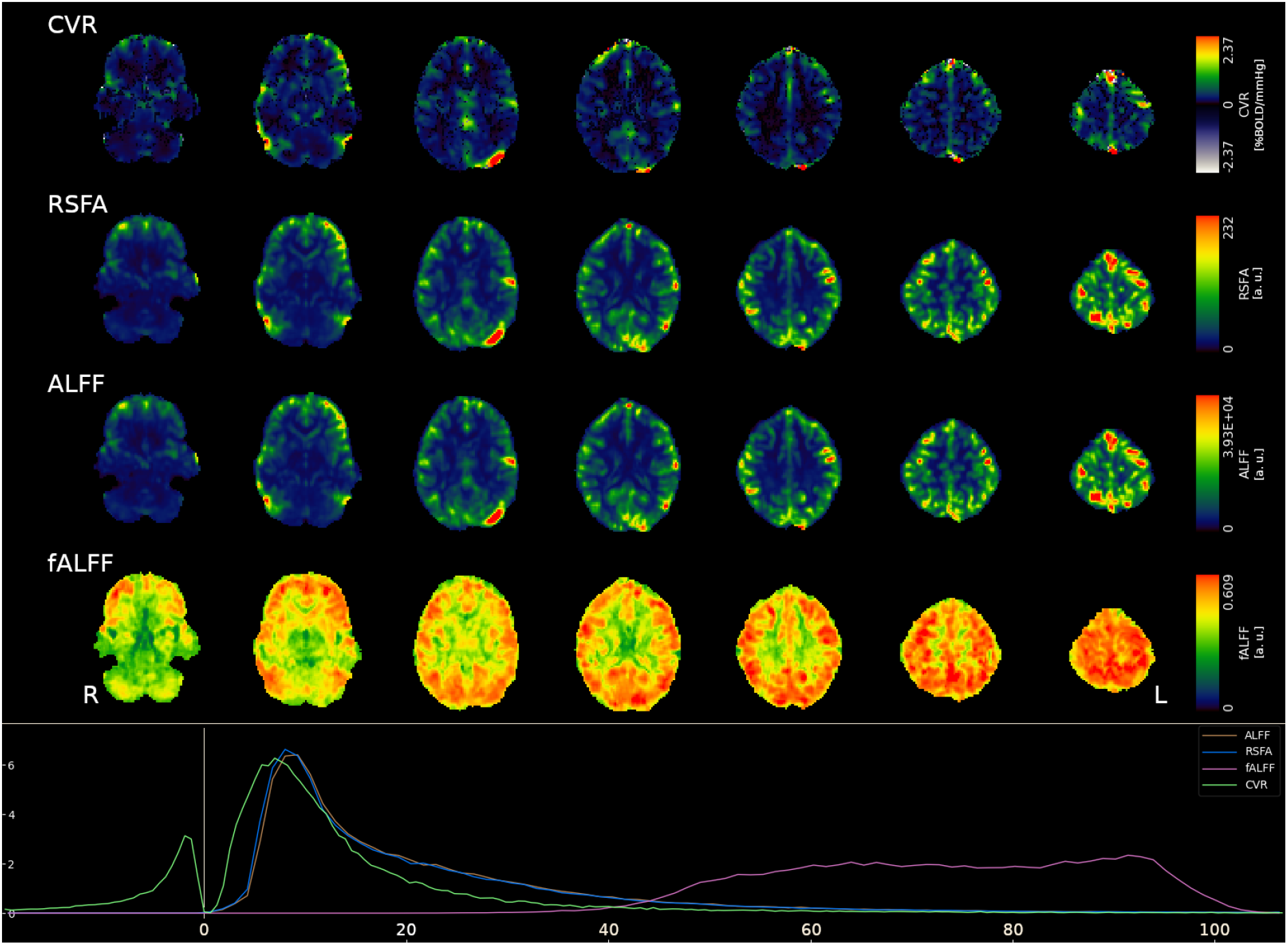
Top: CVR, RSFA, ALFF, and fALFF maps of a representative session of a representative subject. Note that the display range is different for each map, but the upper limit is consistently the 99^th^ percentile of the positive values in the map. CVR is the only metric with negative values. Bottom: Histogram of RSF metrics and CVR values across the whole brain where the *x*-axis represents the percentiles to match their ranges. The map of mALFF is qualitatively equal to the map of ALFF, due to the former being a rescaled version of the latter, as can be seen in Figure S1.

Table 1 indicates the results of the LME modelling of the average GM values of CVR as a function of the RSF metrics, and Figure 2 illustrates these same relationships for ALFF (2A, 2E), RSFA (2B, 2F), fALFF (2C, 2G), and mALFF (2D, 2H). In Figure 2, the left column shows the relationship between CVR and RSF metrics considering all sessions and subjects together, while the right column indicates the relationship for each subject independently. While there seems to be a positive relationship between CVR and RSF metrics when all subjects and sessions are pooled together (i.e. considering all sessions and subjects as independent observations), the statistical probability of rejecting a lack of relationship between CVR and RSF is low, potentially due to the high inter-subject variability revealed by our unique dense mapping dataset. It is also worth noting how, at the group level, the average GM fALFF, i.e. our negative control case, features the second highest fit with average CVR (Figure 2C) compared to RSFA and (m)ALFF. While normally excluded from such comparisons, mALFF shows the largest fit score (*R*2 = 0.98) and the lowest p-value (*p* = 0.11), although again this p-value is not deemed statistically significant.

**Figure 2:**
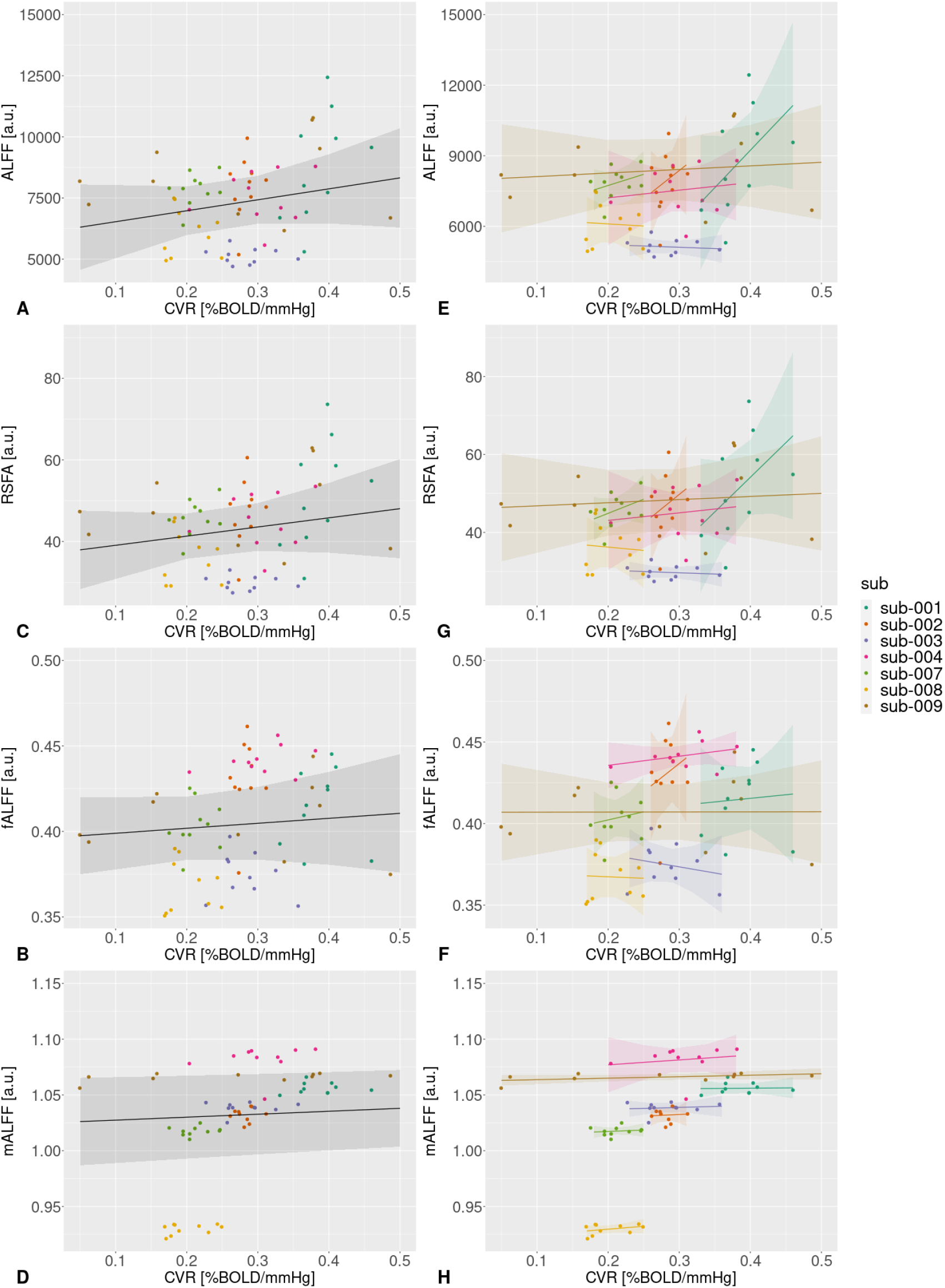
Scatterplots of the relationships between the average GM value of CVR and ALFF (A, E), RSFA (B, F), fALFF (C, G), and fALFF (D, H). The plots in the left column (A-D) illustrate the relationship of all subjects and all sessions pooled together (i.e. considering all subjects and sessions as independent), while the right column (E-H) shows each subject independently. While the left column shows a similar relationship between RSF and CVR compared to previous studies beside statistical significance (cfr. Golestani, Wei & Chen, 2016), the right column shows how this relationship can vary across different subjects.

**Table 1:** LME modelling of average GM RSF metric as a function of the average-GM CVR value for each subject separately and across all subjects (tot). No strong evidence occurred for the presence of relation.

|  | RSFA | ALFF | fALFF | mALFF |
| --- | --- | --- | --- | --- |
| CVR (sub 001) | $\beta = 177.67$<br>$p = 0.16$ | $\beta = 31781$<br>$p = 0.13$ | $\beta = 0.044$<br>$p = 0.85$ | $\beta = 0.005$<br>$p = 0.92$ |
| CVR (sub 002) | $\beta = 147.76$<br>$p = 0.46$ | $\beta = 23933$<br>$p = 0.45$ | $\beta = 0.345$<br>$p = 0.55$ | $\beta = 0.029$<br>$p = 0.85$ |
| CVR (sub 003) | $\beta = -6.45$<br>$p = 0.7$ | $\beta = -1090$<br>$p = 0.73$ | $\beta = -0.075$<br>$p = 0.55$ | $\beta = 0.018$<br>$p = 0.72$ |
| CVR (sub 004) | $\beta = 19.51$<br>$p = 0.7$ | $\beta = 3247$<br>$p = 0.68$ | $\beta = 0.056$<br>$p = 0.32$ | $\beta = 0.044$<br>$p = 0.65$ |
| CVR (sub 007) | $\beta = 72.18$<br>$p = 0.28$ | $\beta = 9503$<br>$p = 0.36$ | $\beta = 0.100$<br>$p = 0.66$ | $\beta = 0.026$<br>$p = 0.68$ |
| CVR (sub 008) | $\beta = -17.08$<br>$p = 0.82$ | $\beta = -1838$<br>$p = 0.87$ | $\beta = -0.018$<br>$p = 0.92$ | $\beta = 0.052$<br>$p = 0.32$ |
| CVR (sub 009) | $\beta = 8.00$<br>$p = 0.74$ | $\beta = 1515$<br>$p = 0.7$ | $\beta = 0.0006$<br>$p = 0.99$ | $\beta = 0.013$<br>$p = 0.13$ |
| CVR (tot) | $\beta = 22.438$<br>$p = 0.41$ | $\beta = 4480.721$<br>$p = 0.34$ | $\beta = 0.029$<br>$p = 0.56$ | $\beta = 0.026$<br>$p = 0.11$ |

Figure 3 depicts the voxelwise results of the model in Equation (1) for RSFA, ALFF, mALFF, and fALFF. The maps are modulated by each voxel’s corresponding z-score, with full opacity corresponding to *p ≤* 0.05 uncorrected. There is a widespread and positive relationship between ALFF and CVR, peaking in visual, parietal, and central lobe, and a negative relationship near the ventricles and in deep white matter, similarly to the results reported by Chen et al. (2023). The relationship between RSFA and CVR exhibits a similar pattern to ALFF, while mALFF features a more extensive negative relationship with CVR than ALFF and RSFA, and fALFF has a strong positive relationship with CVR in the white matter. Note that no voxel survived FDR correction for *p ≤* 0.05. The corresponding thresholded maps are depicted in Figure S2.

**Figure 3:**
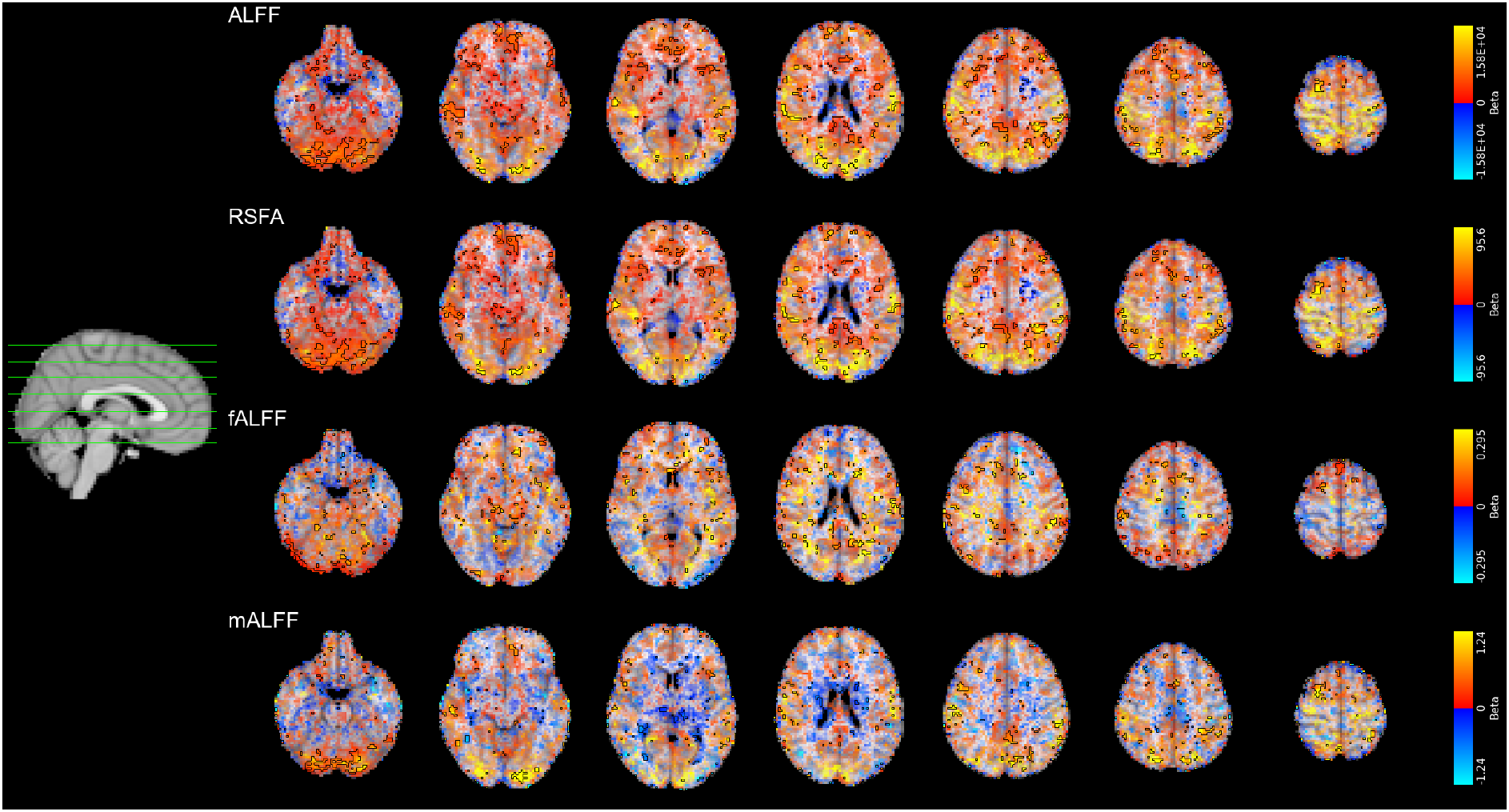
Local (voxelwise) relationship of CVR on RS fluctuations modulated by the maps relative Z score. Areas with *p ≤* 0.05 uncorrected are delimited by black borders. The thresholded version of this figure can be found in supplementary materials (Figure S2).

Figure 4 shows the subject-wise relationship between CVR and RSF metrics for the two most different subjects, while Figures S3, S4, S6, S7, S9, S10, S12, and S13 show the results for all of the subjects (n.b.: the display range is different for each subject to appreciate relative changes in magnitude). The relationship between RSF and CVR differs significantly between subjects, both in size and direction, independently of the metric considered, although it also features certain common patterns. For instance, in Figures S3, S4, S6, and S7 the strongest relationship (relative to each subject) between ALFF/RSFA and CVR seems to be localised in the occipital and in the parietal lobes, although in some subjects these areas feature a positive relationship (cfr. sub-001, sub-007, and sub-009), in others they feature a negative one (cfr. sub-003), and in others this relationship locally shifts from positive to negative, even abruptly (cfr. sub-002, sub-004, and sub-008). Most subjects tend to display a negative relationship between ALFF/RSFA and CVR close to the ventricles, although some feature a positive relationship in the same areas (cfr. sub-002, sub-007, and more prominently sub-008). In the relationship between mALFF and CVR (cfr. Figures S12 and S13), the areas of strongest relationship (relative to each subject) extend to the rostral portion of prefrontal cortex and to the inferior lateral temporal lobes, although once again the sign of such relationship changes between subjects. There seems also to be a prevalence of negative relationship between these two measures, and more subject-specific changes in patterns compared to other RSF measures. Finally, the relationship between fALFF and CVR (cfr. Figures S9 and S10) seems to feature more abrupt changes in sign and magnitude in all subjects, with generally stronger relationship (relative to each subject) in white matter. The subject specificity of these relationships is even more evident when plotting all relationship maps with the same display range (see Figures S5, S8, S11, and S14), since the magnitude of such relationship varies greatly within and between subjects.

**Figure 4:**
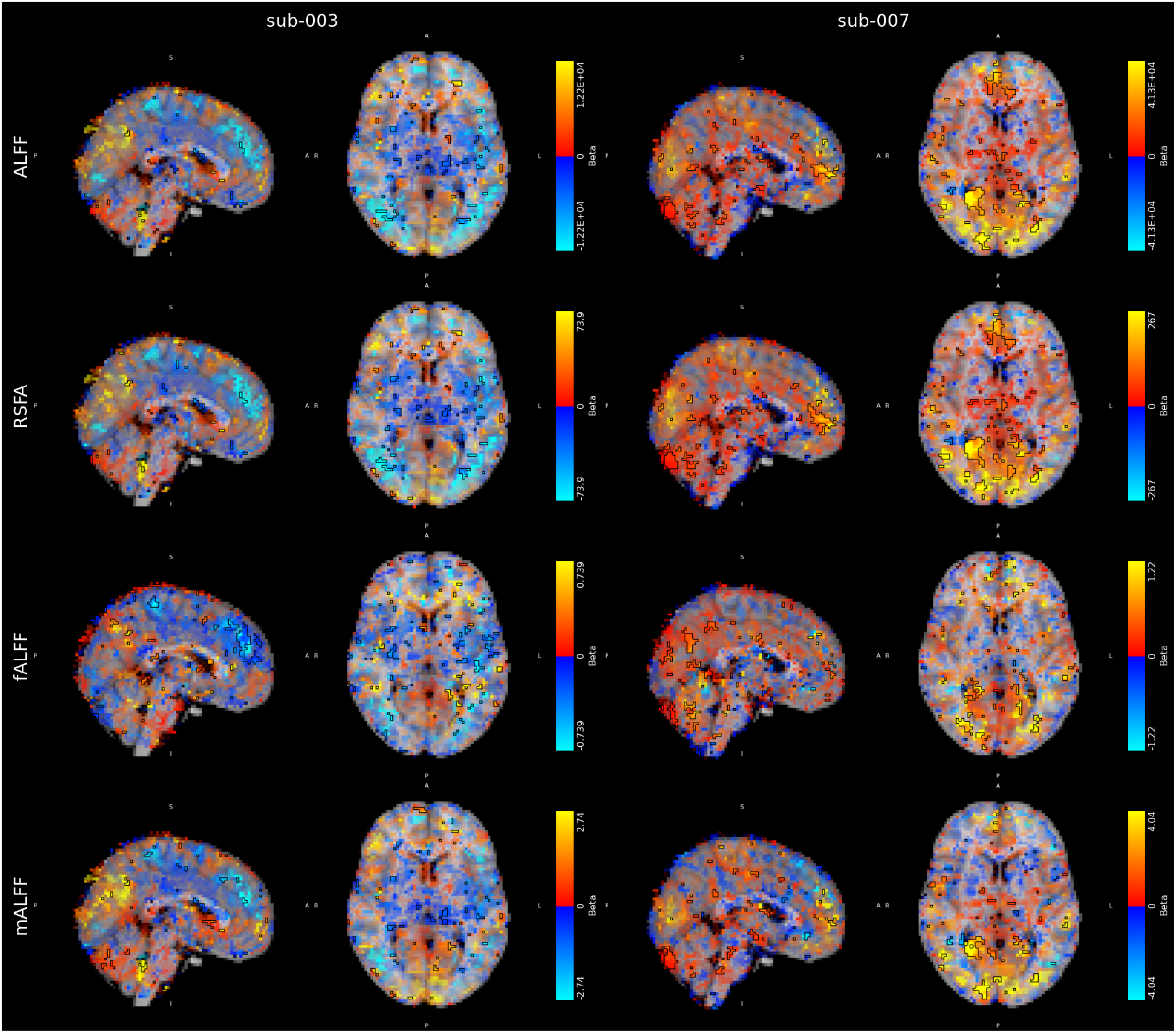
Two examples of subject-wise local relationship between RSF metrics and CVR. Note how the spatial pattern differs between the two subjects, independently of the metric. The effect size is modulated by the relative Z-score (*p ≤* 0.05 uncorrected).

## 4 Discussion

Leveraging the 70 sessions of the first dense mapping fMRI dataset heavily featuring physiological fluctuations, both invoked (i.e. breath-hold) and spontaneous (i.e. resting-state), this study assessed the relationship between RSF metrics (i.e. ALFF, mALFF, fALFF, and RSFA) and CVR (in units of %BOLD/mmHg P_ET_CO_2_), both at the group and at the intra-individual level.

The results revealed that the relationship between CVR and RSF metrics presents high variability across subjects, globally and locally, despite appearing positive at the group level (see Figures 2 to 4). While a good accordance between such measures is found for certain subjects, other subjects do not show this relationship, even featuring a negative relationship in some cases. These observations, only possible thanks to the dense sampling nature of the dataset, demonstrate that between-session (i.e. intra-subject) changes in RSF metrics might not accurately describe between-session (i.e. intra-subject) changes in cerebrovascular physiology. Moreover, they demonstrate the utility that a dense mapping study design can offer for physiological brain imaging.

An easier setup and better patient tolerance of RS fMRI, compared to studies evoking vasodilatory responses using breathing or gas-inhaled challenges, are often cited as reasons to adopt RSF metrics as a proxy for CVR (Fesharaki et al., 2021; Golestani et al., 2016; Kannurpatti & Biswal, 2008; Kannurpatti et al., 2014); in fact, the simplicity of RS fMRI makes it particularly suitable in clinical settings for more vulnerable populations (Fesharaki et al., 2021) and extreme cases such as loss of consciousness (Golestani et al., 2016). Previous studies have even suggested that RSF maps would be better estimators of physiological changes given the high regional agreement between the two measures (De Vis et al., 2018; Kannurpatti & Biswal, 2008; Kannurpatti et al., 2011). As a consequence, RSF metrics have been recognized as potentially more accessible biomarkers of the brain’s vascular health than CVR (Fesharaki et al., 2021; Kannurpatti et al., 2014) due to the lower variability of RSF maps between-subjects (Kannurpatti et al., 2014) or as assessed across the whole brain in a single subject, single session design (Fesharaki et al., 2021). Similar reasons led to the adoption of RSF metrics to rescale the subject-specific BOLD amplitude in task-based activation analysis to account for between-subject variability given the strong similarity between CVR-based and RSF-based rescaling (Di et al., 2013; Kannurpatti & Biswal, 2008; Kannurpatti et al., 2011; Tsvetanov et al., 2015), with the added benefit that RSF metrics can be computed on the same set of data that needs rescaling (Kazan et al., 2016).

However, previous works already demonstrated low spatial (Golestani et al., 2016) and between-subjects (Lipp et al., 2015) agreement between the two measures, and a failure of RSF maps to identify differences between a cohort of internal carotid artery occlusive disease and matched controls (De Vis et al., 2018). Our results, both for voxelwise and average grey matter relationships, show positive, but non-significant, agreement between the two measures at the group level, similar to the work of Lipp et al. (2015), and high variability in the within-subject relationship of RSF and CVR across subjects. While our study includes a limited number of subjects compared to previous literature, the increased number of sessions per subject has a similar impact on statistical efficiency as a bigger sample size, thus allowing such comparisons (Chen et al., 2022). In addition, our results reveal that week-to-week fluctuations in CVR are different from week-to-week fluctuation in RSF. This suggests that either (a) there exists a common driver, such as the BOLD contrast itself, that impacts the two measures, but RSF and CVR are differently impacted over time, or (b) the impact of a common driving property is smaller than originally thought, or should be reconsidered altogether. This interpretation is in line with recent evidence of a stronger link between RSF and haemodynamic responses, specifically neurovascular coupling, than to CVR measured through a BH task (Bailes et al., 2023): while both RSF and CVR might describe a component related to neurovascular coupling, they are not necessarily linked to the same component. Hence, interpreting our results based on our unique dense sampling data, we join the point of view of Lipp et al. (2015) and De Vis et al. (2018) and express caution in adopting RSF measures interchangeably with CVR. For the same reason, we suggest that using RSF metrics for rescaling task activation values might not truly rescale the signal based on vascular changes, rather than on features of the neurovascular coupling and the haemodynamic response (Bailes et al., 2023). Instead, it has also been suggested that the BOLD signal changes induced by the vasodilation associated to breath-holds is strongly influenced by changes in cerebral blood volume (CBV) characterised by deoxy-haemoglobin, and that a novel new marker of deoxygenated CBV with BOLD (BOLD-CBV) can account for more between-subject variability in task-induced BOLD responses (Biondetti et al., 2024).

Note that this interpretation is in direct contrast with that of Kannurpatti et al. (2014), that instead suggested RSF measures to be a better estimator of vascular changes, due to the higher within-subject variability of CVR mapping. However, we have previously shown that our CVR maps are estimated through a procedure to optimise haemodynamic delay while reducing the impact of instantaneous motion (Moia, Stickland et al., 2020), and both BH and RS data are obtained with state-of-the-art MRI acquisitions that improve such estimation (namely, Multi-Echo fMRI, see Kundu et al., 2018). As such, both data types are not only optimised from a denoising perspective, but also highly reliable (Moia et al., 2021). Thus, we do not have evidence that lower variability means that RSF is a better estimator of underlying physiology than BH-induced CVR.

Understanding why there exists such inter-subject variability in the relationship between RSF and CVR requires further study. However, we can proffer a few reasons behind such observations. First, natural respiration and cardiac frequencies can change across subjects, leading not only to different frequency bands describing physiological processes, but also to different aliasing of these effects in the signal (Caballero-Gaudes & Reynolds, 2017). Thus, while we used a fixed and standard frequency band in estimating RSF metrics, a subject-tailored approach might reduce the observed inter-subject variability. Moreover, while the low frequency range is the most adopted range in computing RSF measures, it is important to note that the frequency of events related to vascular responses and their aliases might lie in a smaller or different frequency band. For instance, Liu et al. (2017) suggests adopting the average signal in the 0.02–0.04 Hz bandwidth as the ideal response to vascular fluctuations, in three RS fMRI datasets with TR ranging between 0.27 and 2 seconds. While this work did not consider alternative bandwidths within or outside of the 0.01-0.1 Hz range, future analyses could adopt instead the suggested 0.02-0.04 Hz bandwidth. Alternatively, another option could be considering specific cardiac and respiratory main bandwidths, or their aliases instead.

Second, as previously noted, neuronal activity in RS should be interpreted as "noise" if RSF measurements are to be interpreted as physiological changes, leading to higher variability (Golestani et al., 2016). A limitation of this work is related to the way CVR was obtained. While previous work examining RS and CVR relationships used CO_2_ gas inhalation challenges to obtain CVR (e.g. Golestani et al., 2016), in this work we used a BH challenge instead. While BH-induced CVR shows good agreement to CO_2_-delivery-induced CVR (Kastrup et al., 2001), the required compliance to obtain a good CVR response, and the spurious neuronal effects that a respiratory task likely entails, might influence our results. However, we removed particularly non-compliant subjects, and we conjecture that neuronal effects in the BH task could even increase, and not decrease, the relationship between CVR and RS metrics, due to the task-induced response that characterises neurovascular coupling (Bailes et al., 2023).

It is indisputable that a RS dataset is easier to obtain than a BH dataset. While better RS-based CVR estimation models have been recently proposed, using band-pass filtered signal fluctuations as reference (Liu et al., 2017) or deep learning based algorithms (Hou et al., 2023), recent development in imaging and data analysis methods make BH-induced CVR easier to estimate, and more robust to the lack of subject compliance. While a partial lack of compliance can still lead to reliable CVR estimation (Bright & Murphy, 2013), to reduce even further the necessity of subject compliance, a RS dataset can be slightly altered: for instance, Liu et al. (2020) suggest the adoption of intermitted breath modulation within a RS scan, and Stickland et al. (2021) proposes the addition of a short respiratory challenge before (or after) a period of RS. Both these options can be better tolerated by subjects and introduced in clinical settings, and while relying on RS to increase the amount of analysable data, they gain the benefit of respiratory challenges for improved estimation of CVR. When lack of compliance leads to low quality of collected physiological data, e.g. in the case of poor CO_2_ recordings through nasal cannula, other methods to model CVR response can be used instead (cfr. Golestani et al., 2016; Lipp et al., 2015; Murphy, Harris & Wise, 2011; Pinto, Jorge, Sousa, Vilela & Figueiredo, 2016; Zvolanek et al., 2023) Moreover, improved modelling and denoising can improve CVR estimation as well, for instance by adopting advanced acquisition techniques and lag-based CVR estimations (cfr. Donahue et al., 2016; Moia et al., 2021), deep learning methods to extract CVR (Hou et al., 2023), or, as more recently proposed, algorithms leveraging analyses in the frequency domain, rather than in the time domain (Nanayakkara, Meusel, Anderson & Chen, 2023; Xu et al., 2023).

Conversely, the high inter-subject variability and frequent absence of a relationship between CVR and RSF metrics could be related to how evoked and spontaneous CO_2_ fluctuations might produce vascular responses that differ in magnitude and temporal definition (i.e. time-locked or spontaneous). Indeed, previous comparisons of CVR estimated in RS or during a gas challenge showed a few differences between the two, notably the presence of an undershoot in spontaneous CVR (and the lack of one in induced CVR), and a small difference in CVR magnitude (Prokopiou, Pattinson, Wise & Mitsis, 2019). While previous studies reporting a modest relationship between CVR and RSF suggest to interpret CVR and RSF as two different aspects of vascular changes, only partially sharing similar characteristics (Bailes et al., 2023; Chen et al., 2023; Lipp et al., 2015; Liu et al., 2013), further studies addressing specific aspects of vascular responses to different CO_2_ stimuli (or the lack of) are required. Yet, using CVR estimates obtained from CO_2_ recordings during the RS run could be an alternative approach to using BH-induced CVR estimates in specific pathological cases where compliance to a task is impossible to obtain, e.g. in the case of lack of consciousness (Golestani et al., 2016). In such cases, the main limitation in estimating CVR from RS data is that estimating the voxelwise haemodynamic delay is more challenging than computing RSFA, ALFF, mALFF, or fALFF when lacking a CVR inducing task (Stickland et al., 2021): it is more difficult to individuate true peaks in the CO_2_ signal during normal (or mechanically ventilated) breathing, and the fMRI signal during spontaneous fluctuations exhibits lower variability (i.e. contrast-to-noise ratio) than in BH data. Further development of alternative modelling strategies, such as those based in frequency domain (Nanayakkara et al., 2023; Xu et al., 2023) or on deep learning (Hou et al., 2023) might improve the specificity of RS fMRI estimates of CVR, and reduce the inter-subject variability revealed in the current study.

Finally, to our knowledge this is the first application of a dense mapping fMRI dataset acquisition focusing on cerebrovascular physiology. Dense mapping datasets are characterised by acquiring multiple sessions, extending traditional test-retest studies, in a few subjects, and have been gaining popularity in recent years. This popularity came from pioneering studies in the neuroimaging field showing how dense mapping can bring new insight on the impact of external factors on functional connectivity (Poldrack et al., 2015), on peculiarities of functional subnetworks (Braga & Buckner, 2017), or on changes over time in functional connectivity (Laumann et al., 2015) and network dynamics and interactions (Gordon et al., 2018; Gratton et al., 2018). More recently, dense mapping has been used both in cognitive and neurobiological studies, for instance observing the effect of brain plasticity (Newbold & Dosenbach, 2021; Newbold et al., 2020) and functional reconfiguration (Salehi, Karbasi, Barron, Scheinost & Constable, 2020), or the effect of endocrine system during menstrual cycle (Pritschet et al., 2020; Pritschet, Taylor, Santander & Jacobs, 2021), as well as in clinical settings, for instance comparing the functional connectivity of individuals with depression against that of healthy individuals (Lynch et al., 2023). In the current study, dense mapping allowed us to assess how the inter-session relationship between RS metrics and CVR changes across individuals, thus showing the exciting potential of this acquisition approach in investigating cerebrovascular physiology with fMRI from a different perspective.

## 5 Conclusion

This study investigated the relationship between CVR induced by a BH task and RS fluctuations in a dense mapping fMRI dataset with 7 subjects each scanned during 10 sessions, considering such relationships at both the group and at the subject level. We showed a similar cross-sectional relationship between RSF metrics and CVR to previous studies (e.g. Golestani et al., 2016), although in our dataset such relationship was not found to be significant. However, when looking into the same relationship at the subject level, we showed high variability between subjects.

We also found no significant widespread relationship when considering RSF metrics and CVR at the voxelwise level. These results suggest caution in interpreting RSF metrics as markers of vascular physiology akin to CVR, or to use them to calibrate subjects fMRI responses based on physiological fluctuations.

Such new insights in a highly debated topic are possible only due to the dense mapping study design, which opens up new research avenues into cerebrovascular function.

## S1 Supplementary materials

**Figure S1:**
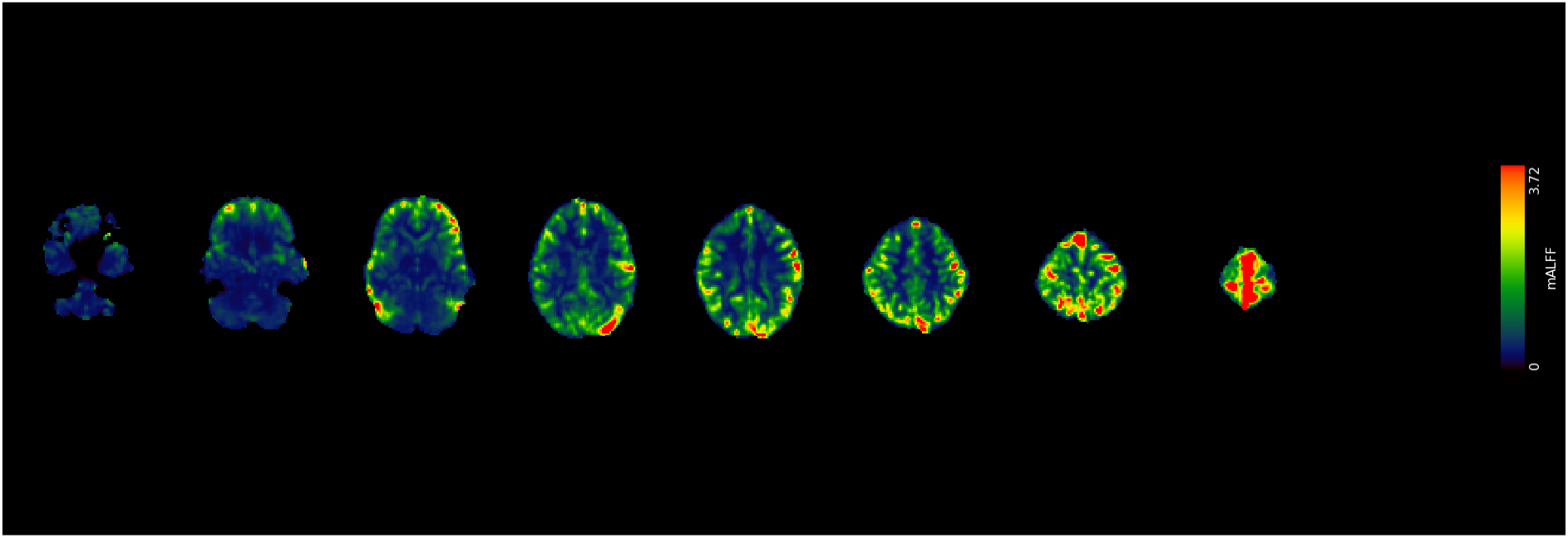
mALFF maps of a representative session of a representative subject.

**Figure S2:**
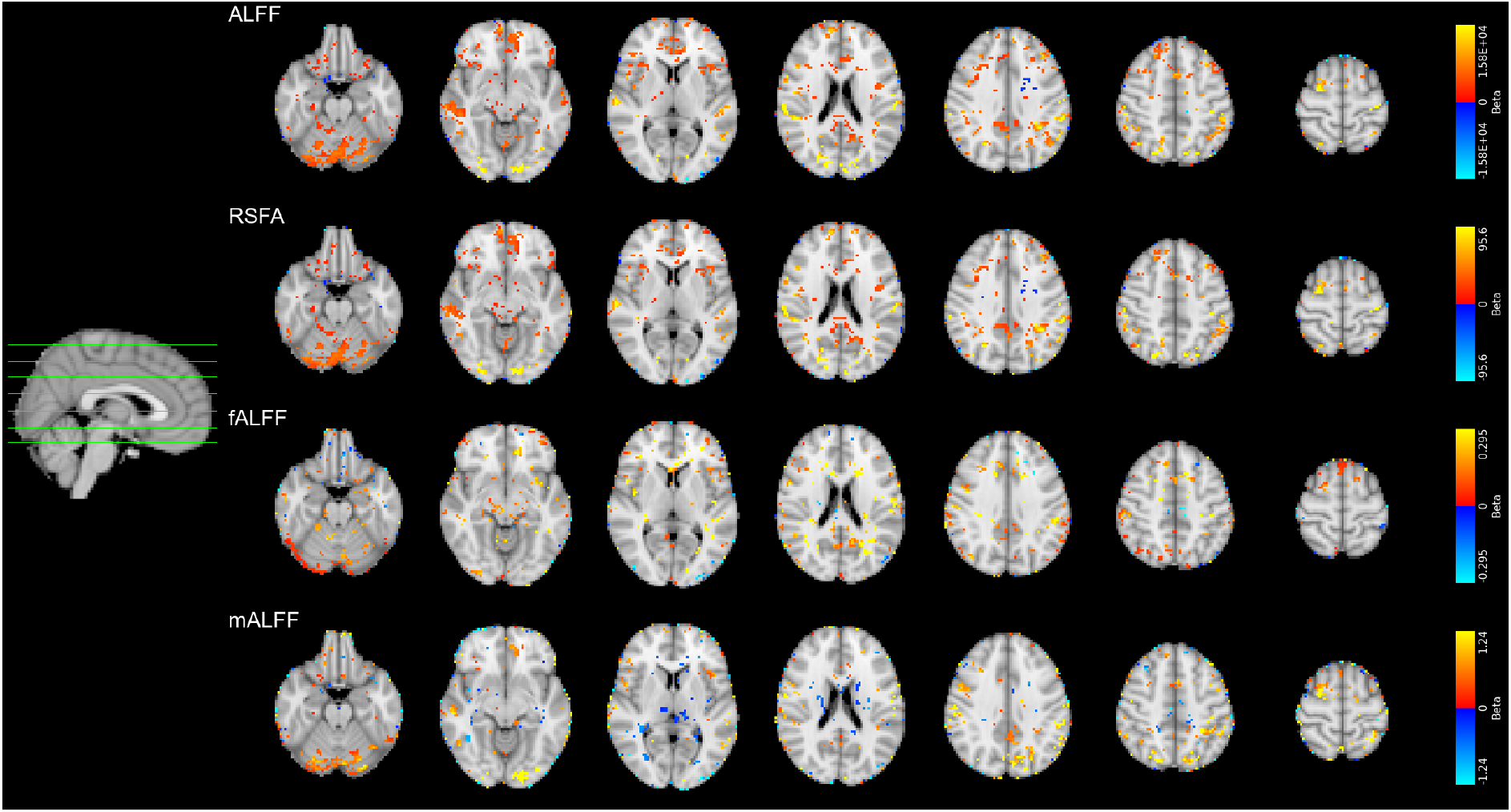
Local effect of CVR on RS fluctuations (*p ≤* 0.05 uncorrected).

**Figure S3:**
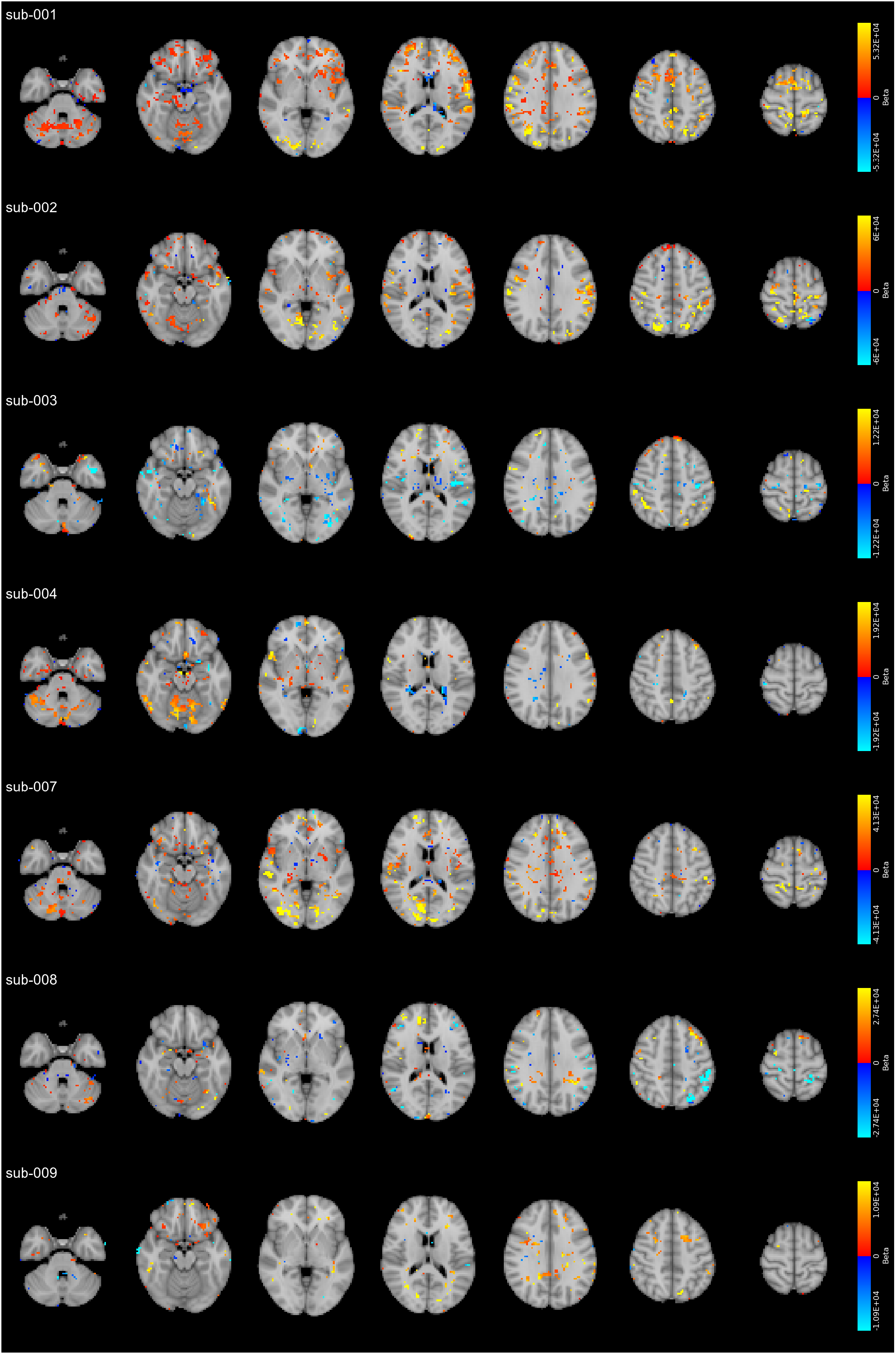
Local effects of CVR on ALFF for each individual subject, thresholded at *p ≤* 0.05 uncorrected

**Figure S4:**
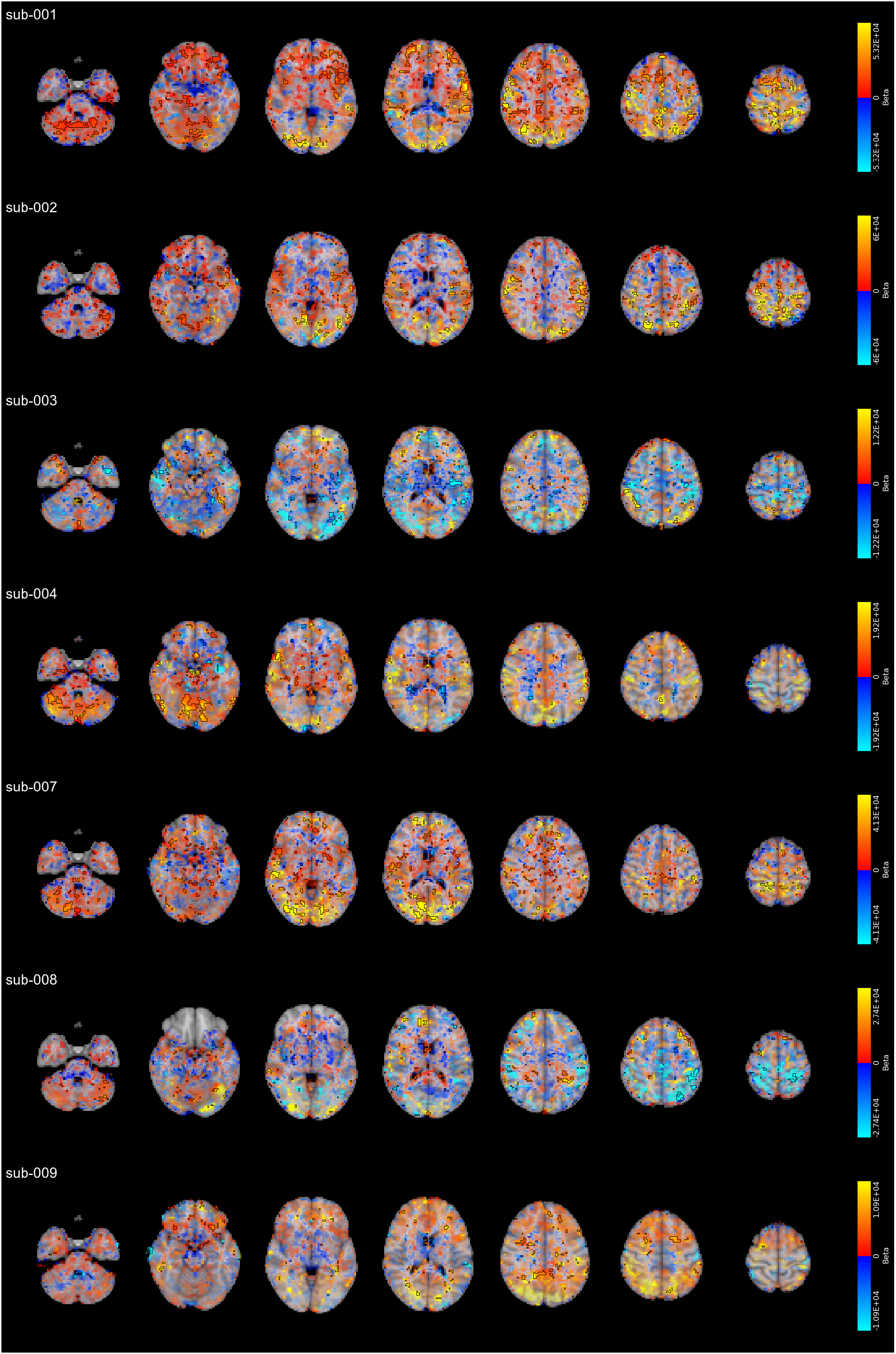
Local effects of CVR on ALFF for each individual subject, modulated by the maps relative Z score (*p ≤* 0.05 uncorrected), featuring a different display range for each subject (98th percentile of the relationship strength distributions)

**Figure S5:**
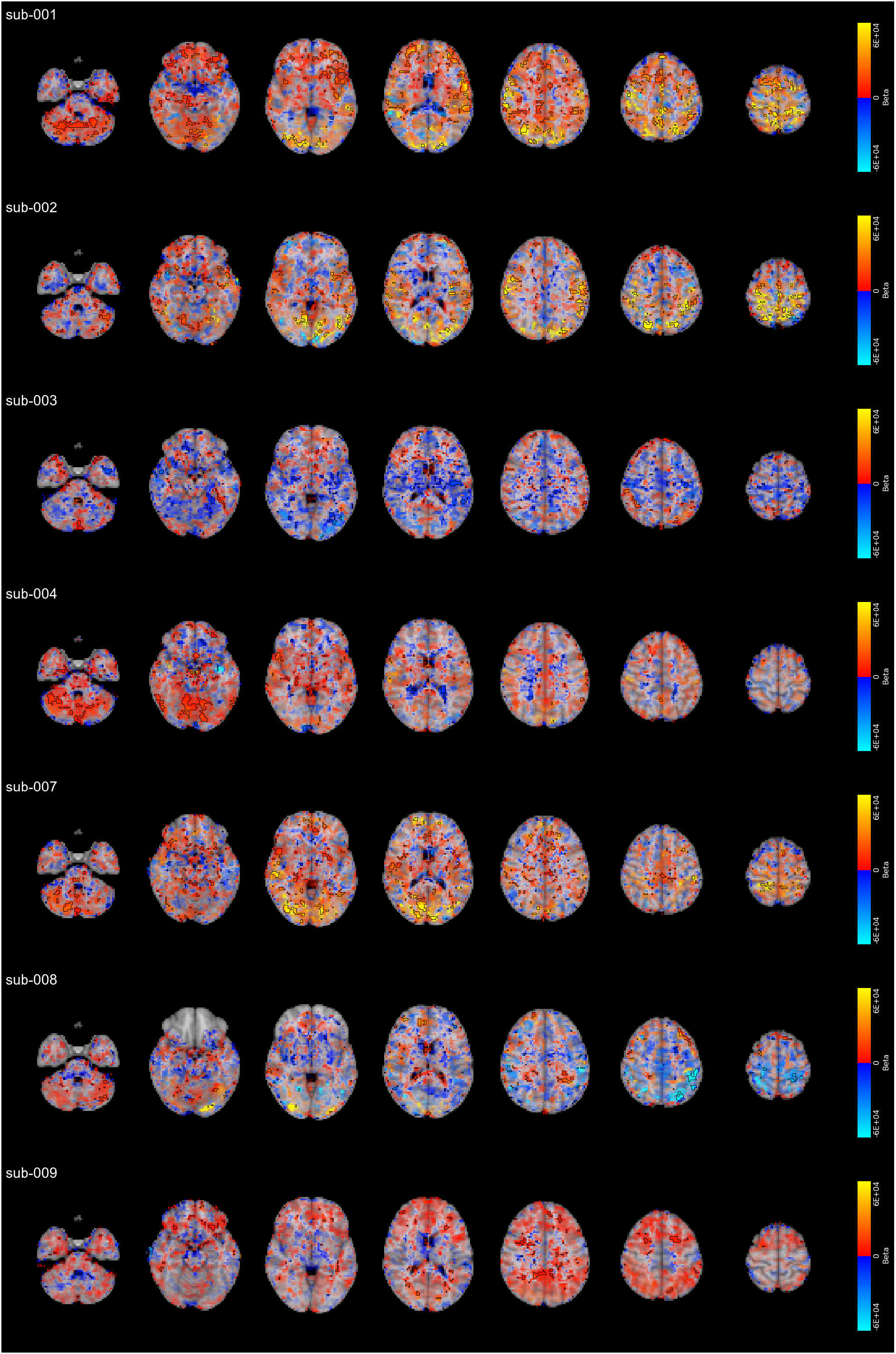
Local effects of CVR on ALFF for each individual subject, modulated by the maps relative Z score (*p ≤* 0.05 uncorrected), featuring the same display range for all subjects (based on the 98th percentile of the relationship strength distribution of the subject with strongest relationship)

**Figure S6:**
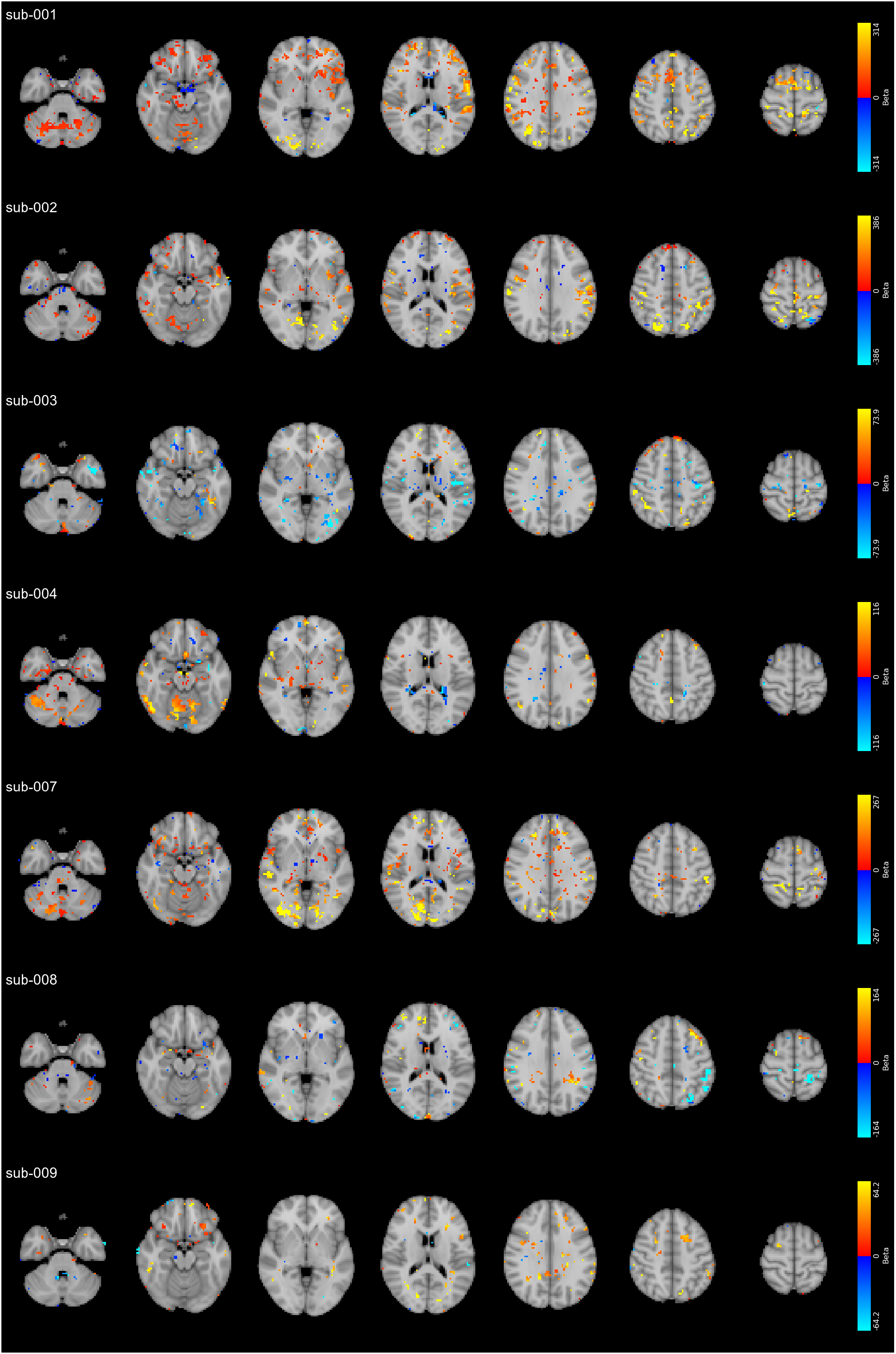
Local effects of CVR on RSFA for each individual subject, thresholded at *p ≤* 0.05 uncorrected

**Figure S7:**
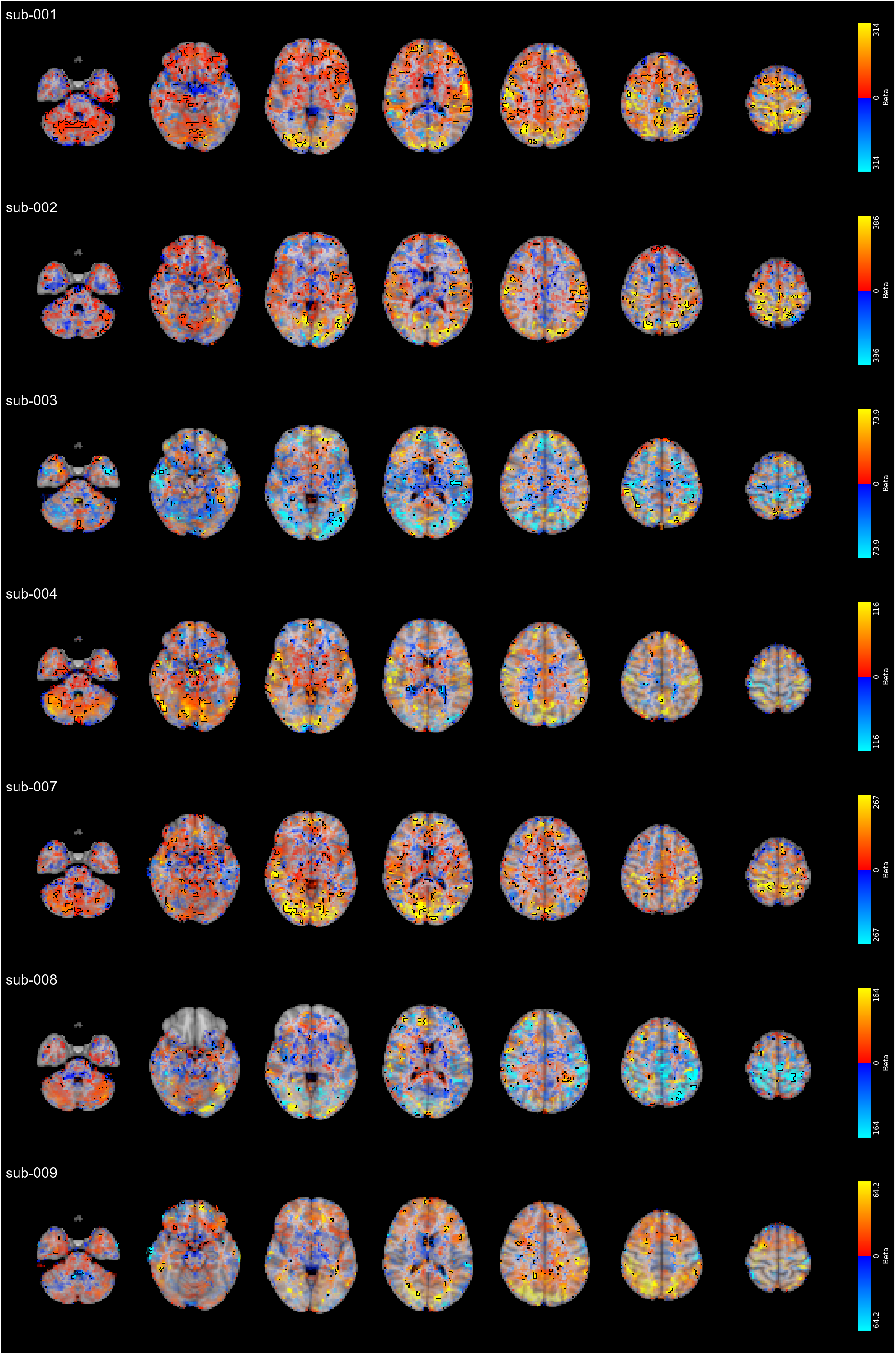
Local effects of CVR on RSFA for each individual subject, modulated by the maps relative Z score (*p ≤* 0.05 uncorrected), featuring a different display range for each subject (98th percentile of the relationship strength distributions)

**Figure S8:**
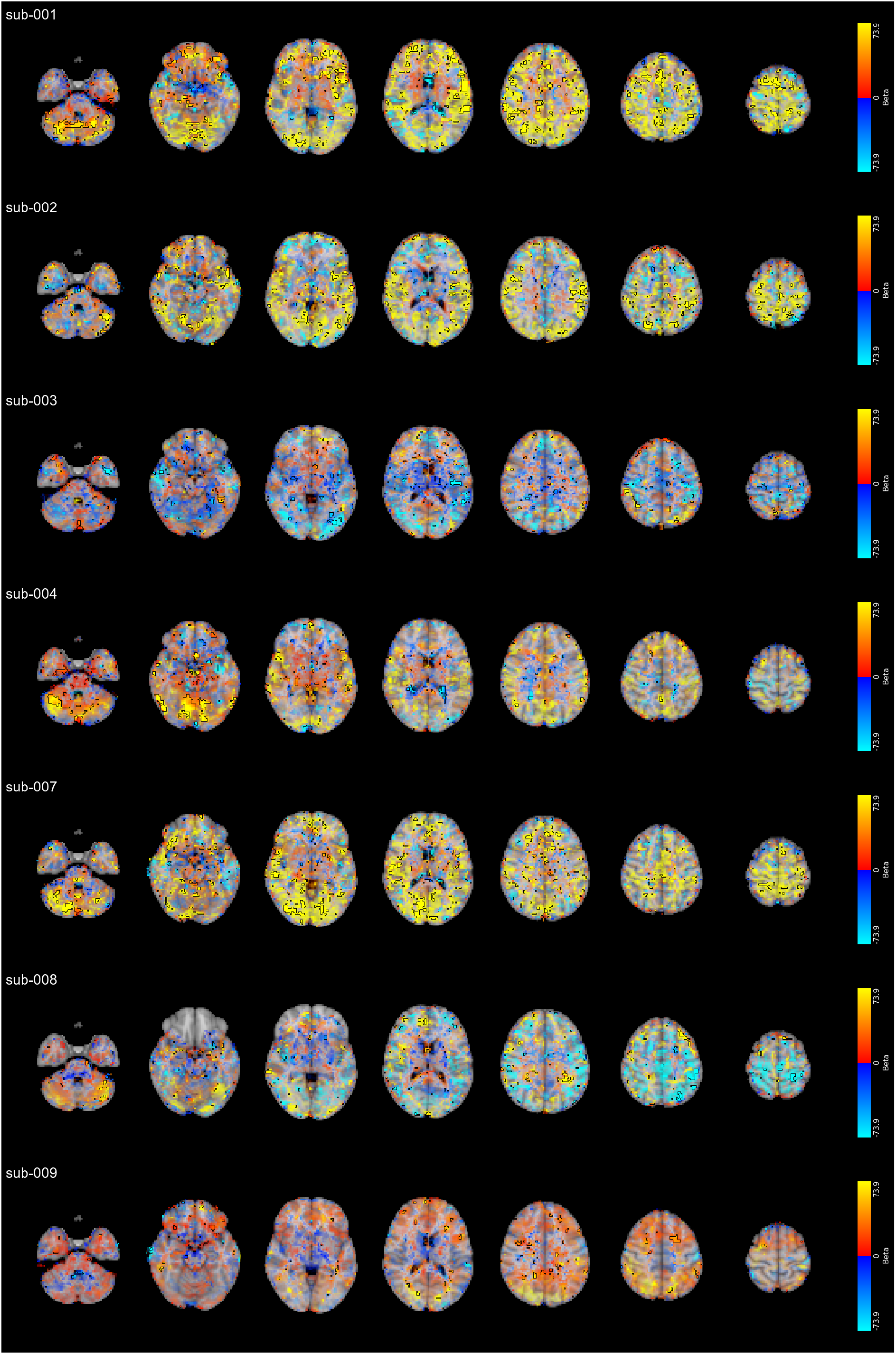
Local effects of CVR on RSFA for each individual subject, modulated by the maps relative Z score (*p ≤* 0.05 uncorrected), featuring the same display range for all subjects (based on the 98th percentile of the relationship strength distribution of the subject with strongest relationship)

**Figure S9:**
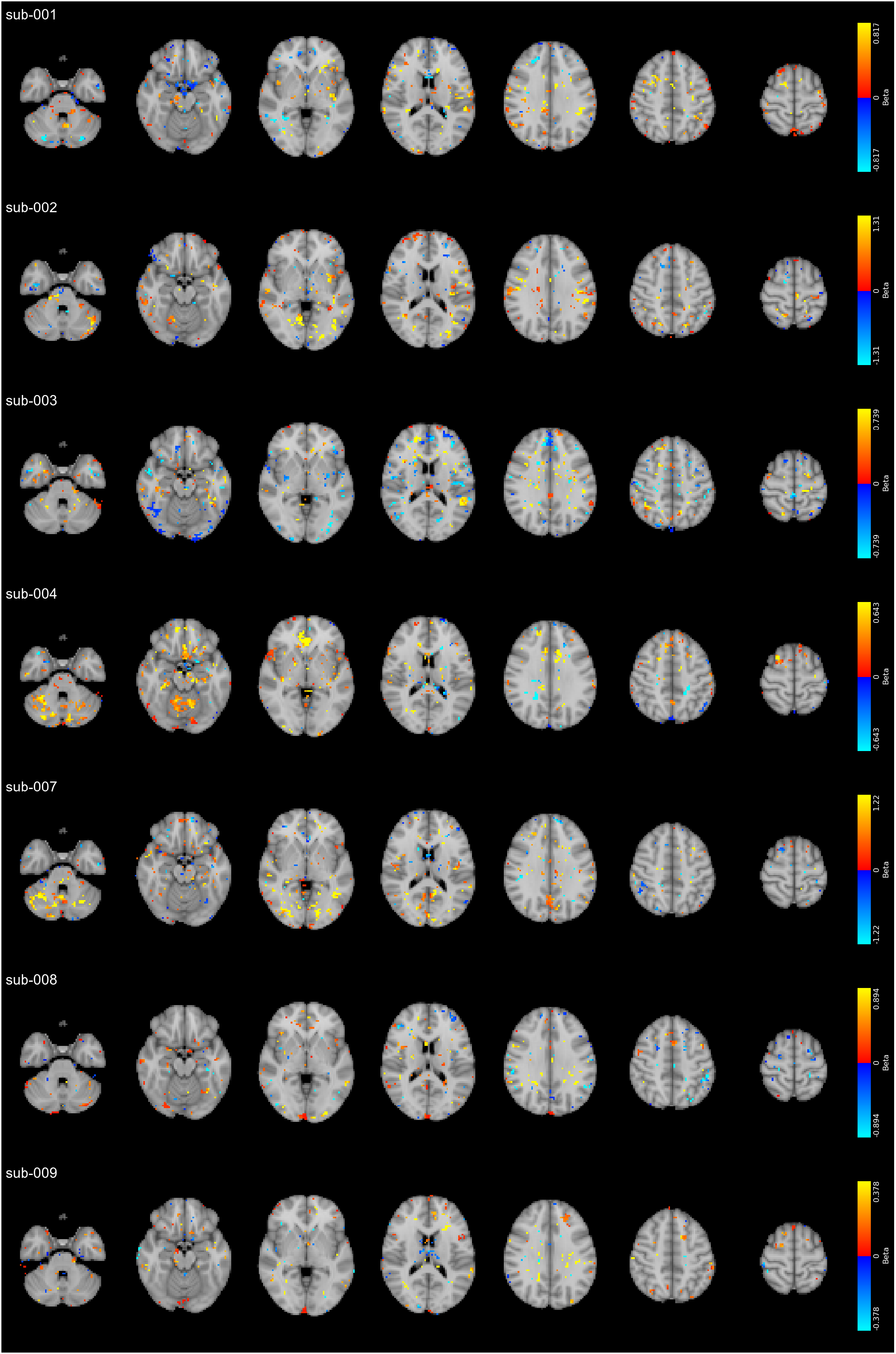
Local effects of CVR on fALFF for each individual subject, thresholded at *p ≤* 0.05 uncorrected

**Figure S10:**
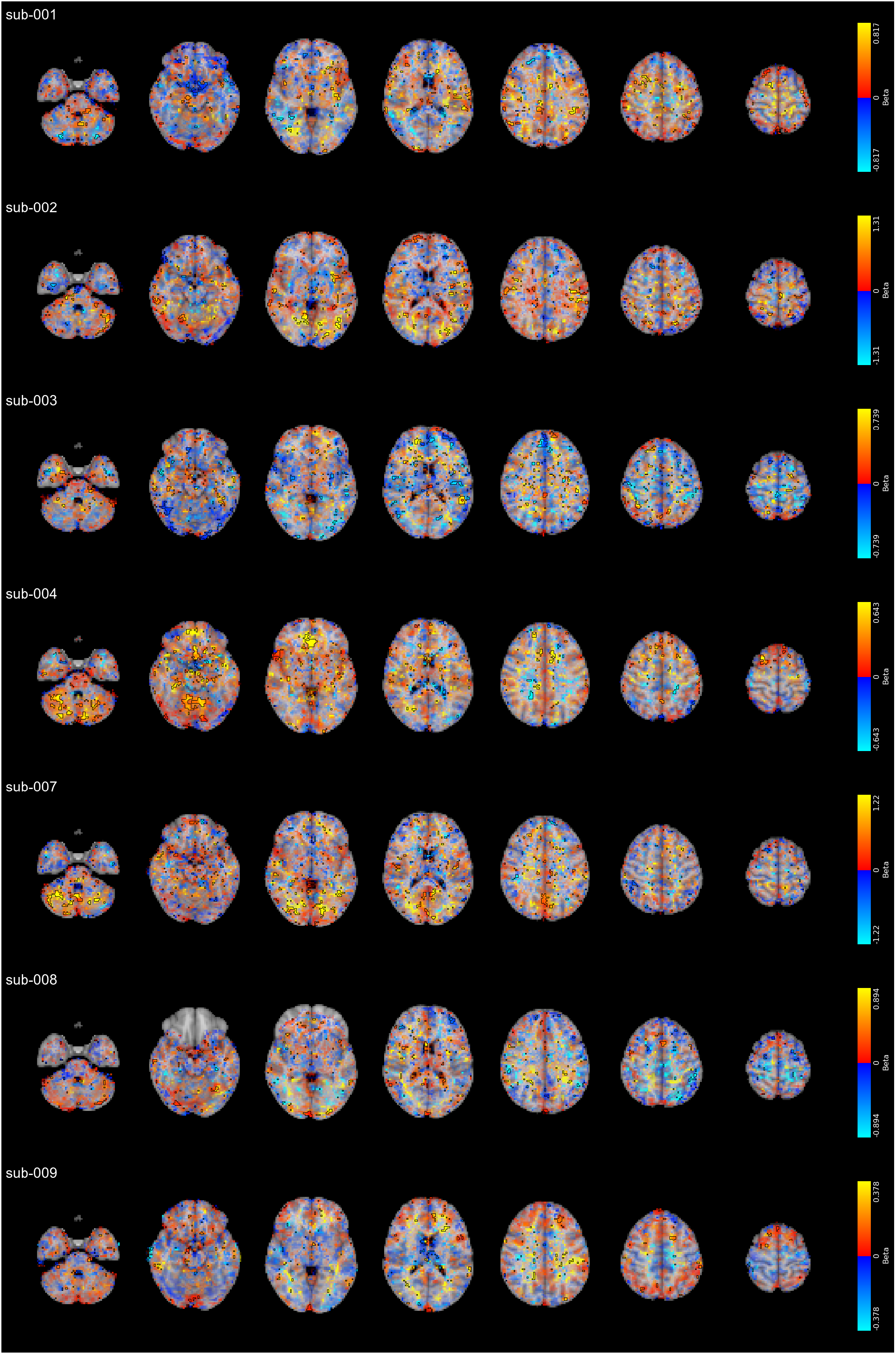
Local effects of CVR on fALFF for each individual subject, modulated by the maps relative Z score (*p ≤* 0.05 uncorrected), featuring a different display range for each subject (98th percentile of the relationship strength distributions)

**Figure S11:**
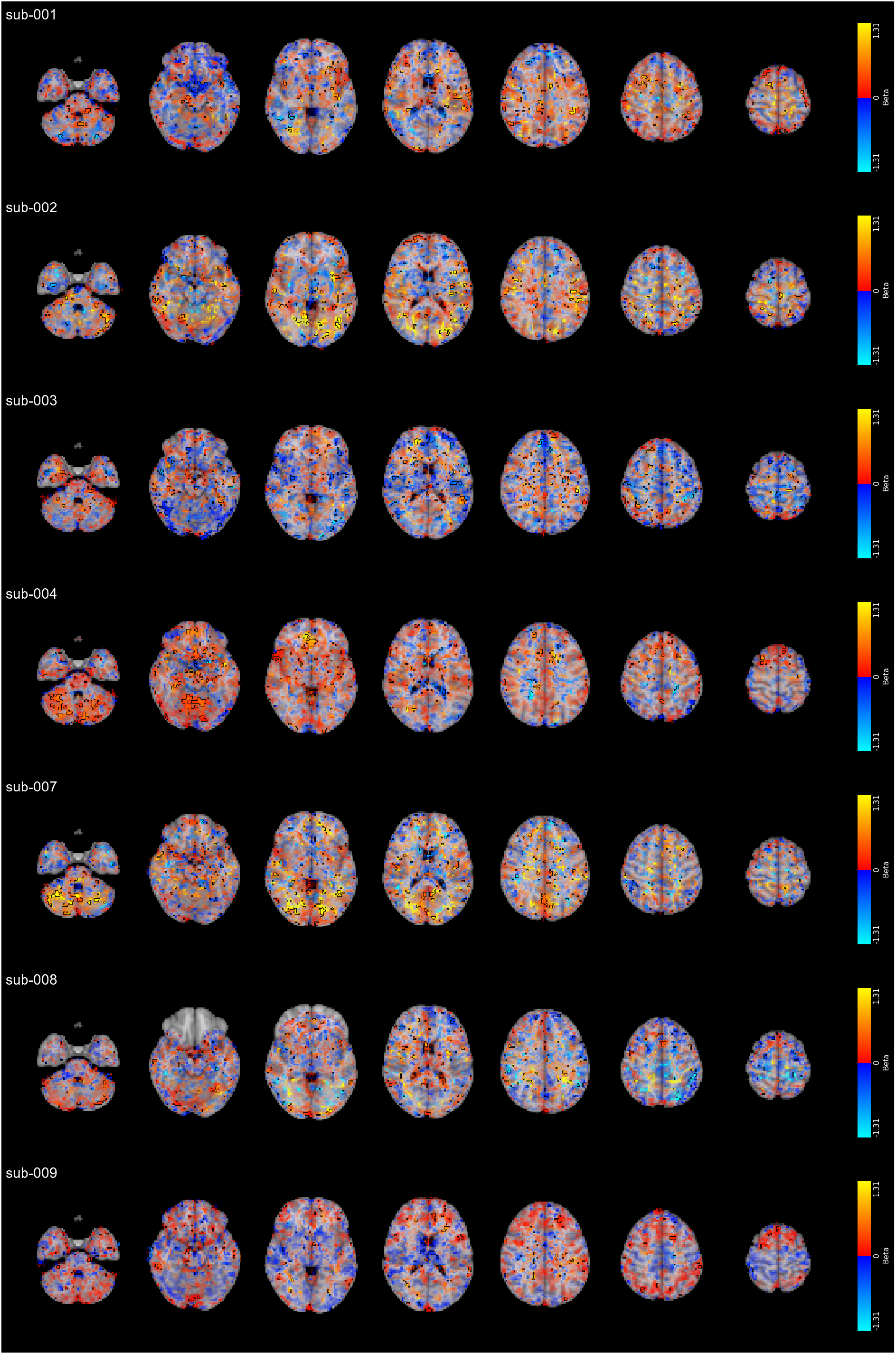
Local effects of CVR on fALFF for each individual subject, modulated by the maps relative Z score (*p ≤* 0.05 uncorrected), featuring the same display range for all subjects (based on the 98th percentile of the relationship strength distribution of the subject with strongest relationship)

**Figure S12:**
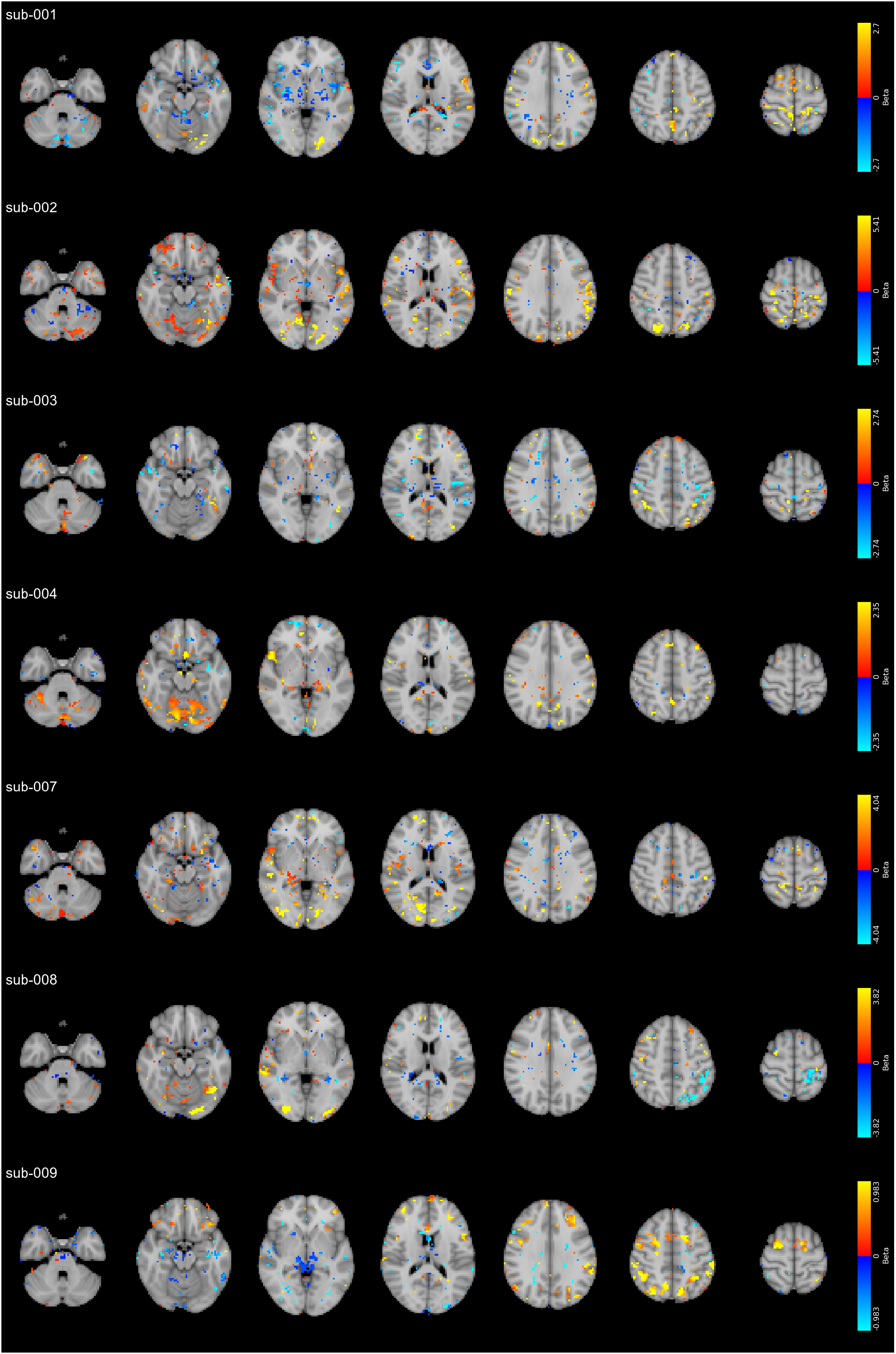
Local effects of CVR on mALFF for each individual subject, thresholded at *p ≤* 0.05 uncorrected

**Figure S13:**
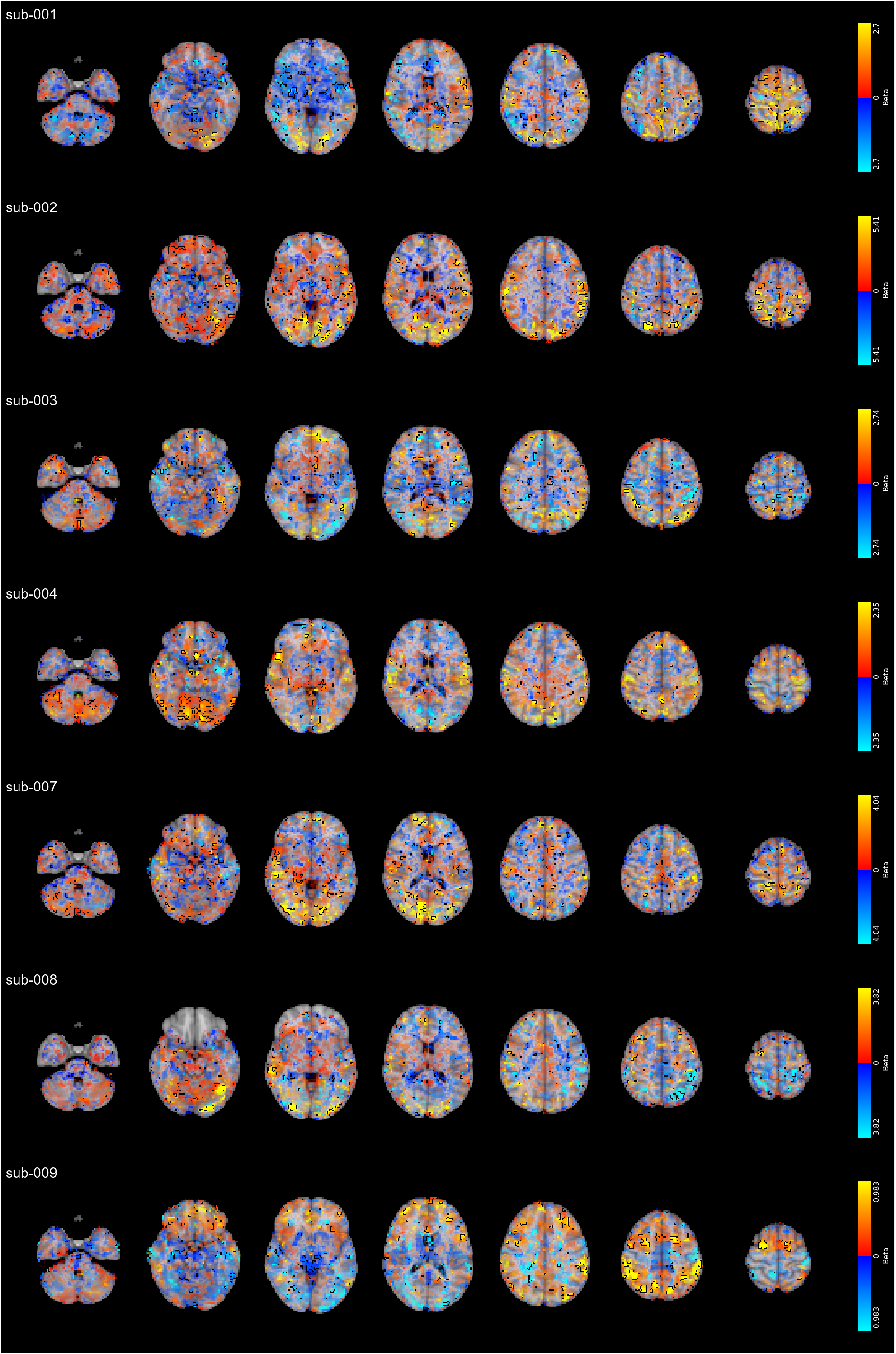
Local effects of CVR on mALFF for each individual subject, modulated by the maps relative Z score (*p ≤* 0.05 uncorrected), featuring a different display range for each subject (98th percentile of the relationship strength distributions)

**Figure S14:**
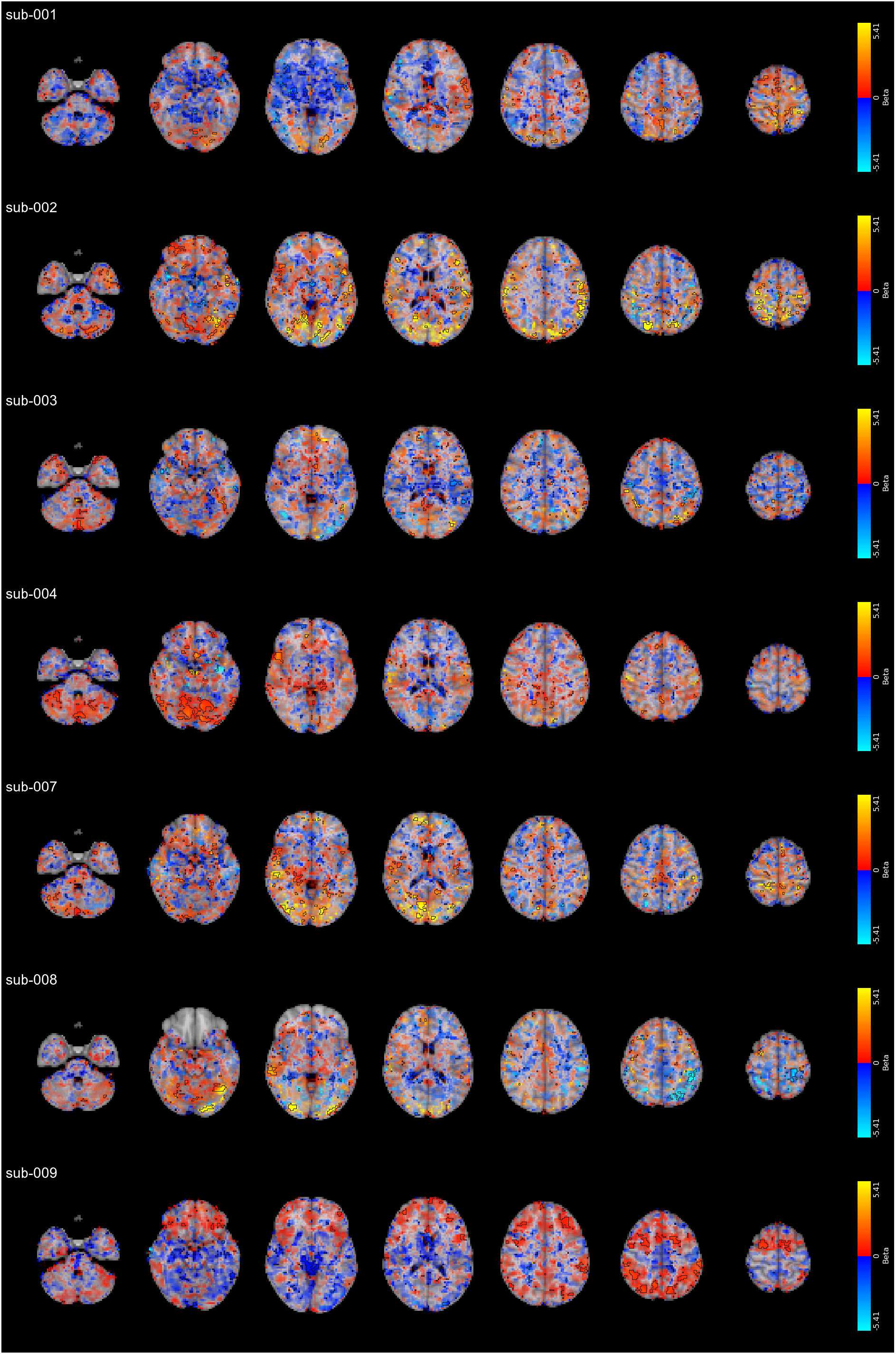
Local effects of CVR on mALFF for each individual subject, modulated by the maps relative Z score (*p ≤* 0.05 uncorrected), featuring the same display range for all subjects (based on the 98th percentile of the relationship strength distribution of the subject with strongest relationship)

## Footnotes

1 https://github.com/smoia/EuskalIBUR_dataproc

2 https://git.bcbl.eu/smoia/euskalibur_container

